# FiberTron enables regulatory variant effect prediction and testing from multi-omic long reads of a single sample

**DOI:** 10.64898/2026.11.13.762587

**Authors:** Marios Eftychiou, Laurens Lambrechts, Joris Vande Velde, Koen Theunis, Christopher Davies, Nischalan Pillay, Kim De Keersmaecker, Stein Aerts, Alejandro Sifrim, Jonas Demeulemeester

## Abstract

State-of-the-art sequence-to-function models are trained on large compendia of public data to predict regulatory activity across hundreds of cell types. However, they are not designed to interpret individual genomes, effects in under-sampled cell types, or provide a comprehensive view of how genetic variation influences complex diseases. Here we show that a single sample profiled with Fiber-seq and Nanopore direct RNA-seq contains enough information to train such a model from scratch. FiberTron, a 12M parameter convolutional–transformer model, takes 524 kb of reference sequence and predicts CpG methylation, chromatin accessibility and strand-specific transcript coverage at base-pair resolution (median Pearson’s *r* of 0.85–0.97 on held-out chromosomes). Trained in under a day on a single four-GPU node, our HG002 model performs on par with models such as AlphaGenome on dsQTL and eQTL variant effect prediction tasks, despite being over an order of magnitude smaller. As training uses reference sequence only, each sample’s phased multi-omic long reads remain available to test its variant-effect predictions. FiberTron recapitulates *OAS1* splicing variation associated with COVID-19 risk, recovers 79% of allele-specific accessibility variants in a melanoma line, identifies oncogenic *TERT* promoter and *TAL1* enhancer mutations, and predicts *EWSR1::ATF1* fusion transcript expression and its unique chromatin imprint in a rare clear cell sarcoma patient sample. Lightweight single-sample training brings base-resolution interpretation of non-coding variation within reach of individual laboratories.

## Introduction

Understanding the functional impact of non-coding genetic variation has been a long-standing goal in the field of genomics. Over 98% of observed genetic variation in the human genome is non-coding^1^. In conjunction with the diverse and sometimes complex molecular effects non-coding variants can elicit^2^, this makes the interpretation of their impact intractable without extensive wet-lab experiments or computational methods.

The successful adoption of state-of-the-art (SoTA) Deep Learning (DL) methods from the fields of signal-processing and computer vision has led to an explosion of studies using DNA sequence to predict high-dimensional functional genomic modalities (i.e., transcription factor binding, chromatin accessibility, gene expression, etc.), in an attempt to investigate and explain the effect of genetic variation on genome function^3^. DL architectures to study the effect of genetic variation mainly fall into two categories; lightweight models specializing in one or few modalities from a single tissue/cell-type such as BPNet^4^, ChromBPNet^5^ and DeepMEL2^6,7^ and heavyweight foundational models capturing diverse functional genomic modalities across multiple species/tissues/cell-types like Enformer^8^, Borzoi^9^ and more recently AlphaGenome^10^.

There is a trade-off between the two categories^3^. Lightweight specialized models tend to be simpler in architecture and cheaper computationally. However, they use limited genomic context input, usually spanning only a few kilobases (kb), which does not allow them to capture the effects of distal regulatory elements. In contrast, heavyweight foundational models can take expanded genomic sequences ranging from several hundred kilobases up to a megabase at the expense of computational complexity. To the best of our knowledge, there are no hybrid models in-between these two extremes, i.e., lightweight single-class (tissue or cell-type) multi-modal models able to handle expanded genomic sequence inputs and capture long-range gene regulatory effects.

This phenomenon may, at least in part, reflect the specific types of functional genomics data employed in prior studies (e.g., DNase-seq, ChIP-seq, ATAC-seq, etc.). While these functional genomic assays have revolutionized our understanding of biology, they have inherent limitations. By design, these count- or peak-based short-read techniques do not allow reconstruction of the original biological macromolecules or their states and therefore provide only partial representations of the genome, epigenome, and transcriptome^11^. Consequently, information may be implicitly excluded or underrepresented during model training. Recent innovations in third-generation sequencing technologies^11^ (Oxford Nanopore Technologies (ONT) and Pacific Biosciences (PacBio) HiFi sequencing) have accelerated our understanding of both genetic and epigenetic variation present in the human genome^12^. These single-molecule long-read sequencing approaches improve genome assembly, mapping, phasing, and identification of structural variants and transcript isoforms^13^. Moreover, the ability to read native DNA molecules without amplification enables direct detection of base modifications.

Methods such as Fiber-seq^14^ and ONT’s direct RNA-seq^15^ leverage this capability to provide base-pair resolution single-molecule multi-omic information. Fiber-seq simultaneously captures long-range sequence with phased, small and complex, genetic variation, DNA methylation (endogenous 5-methylcytosine, 5mC) and open chromatin information (exogenously labeled N6-methyladenine, 6mA). ONT’s direct RNA-seq method allows direct sequencing of RNA molecules and can provide base resolution information for full-length transcript isoforms as well as native RNA modifications. These technological advancements allow high-throughput direct linking of genotypes to molecular phenotypes at single-molecule level and boost our understanding of the role of both small and complex genetic variation in health and disease^14–18^. Moreover, the synergy between long-read multi-omic sequencing and DL is establishing a powerful new paradigm for investigating genome regulation, providing predictive insights into chromatin architecture, gene regulation, and the molecular basis of human disease^19,20^.

Recent advances in S2F modeling have largely been driven by scaling both training datasets and model architectures^9,10,21^. Although these approaches have achieved impressive predictive performance, their development and application typically require extensive computational resources and large consortium-generated datasets. Furthermore, while aforementioned S2F applications capture specific facets of *cis*-regulatory biology, they are still hindered by their inability to reliably interpret individual genomes^22,23^, to quantify regulatory effects in rare understudied cell types^24^, and to provide a broader characterization of how genetic variants influence complex diseases^24,25^.

To this end, we developed FiberTron, a hybrid convolutional neural network architecture that integrates a hierarchical self-attention mechanism to model and interpret information-dense Fiber-seq and direct RNA-seq data. Individual FiberTron instances were trained on whole-genome and whole-transcriptome long-read single-molecule multi-omic data derived from single samples and enable accurate, base-pair resolution prediction of genome-wide DNA methylation, chromatin accessibility, and strand-specific gene expression. Although trained on single-sample data, FiberTron’s predictive performance is on par with substantially larger compendium-trained SoTA foundational models across various variant effect prediction tasks. This extends sequence-based regulatory interpretation to biological contexts that remain underrepresented in public resources, including rare cell types, tumors, and individual patients. In addition, FiberTron’s single-sample training framework avoids extensive training or fine-tuning of heavyweight models, thereby bringing S2F modeling within reach of individual laboratories, while its in-sample validation reduces the need for further experimental verification of phased candidate *cis*-regulatory variants. Finally, by coupling FiberTron’s predictions with explainable AI approaches, trained model instances delineate *cis*-regulatory consequences of non-coding variants in both physiological and disease-associated states.

## Results

### A lightweight single-class multi-modal sequence-to-function model

FiberTron is a S2F DL architecture designed to learn and decipher the genetic regulatory code of the specific biological context it is trained on (i.e., tissue type, cell line, cell type, health/disease phenotype, etc.). By taking DNA sequences of 524 kb size as input, it predicts simultaneously (epi)genomic modalities at base resolution including DNA methylation and chromatin accessibility (Fiber-seq) as well as strand-specific gene expression coverage (direct RNA-seq).

FiberTron employs a hybrid convolutional transformer architecture. It has a U-Net inspired backbone architecture^9,26^ with a hierarchical self-attention mechanism to efficiently process input DNA sequences into multi-resolution-level representations (Figure 1a, Extended Data Figure 1a-b). The convolution layers serve as flexible non-linear pattern detectors with early layers extracting local sequence patterns (e.g., transcription factor binding motifs) and subsequent layers capturing higher-order syntax rules without predefined features^4^. The interleaved transformer modules with self-attention mechanism allow the exchange of information between every pair of these progressively coarser features. Therefore, incorporating these components may enable the model to represent long-range dependencies in the input DNA sequences corresponding to distal interactions between regulatory elements (i.e., enhancers, promoters).

**Figure 1.**
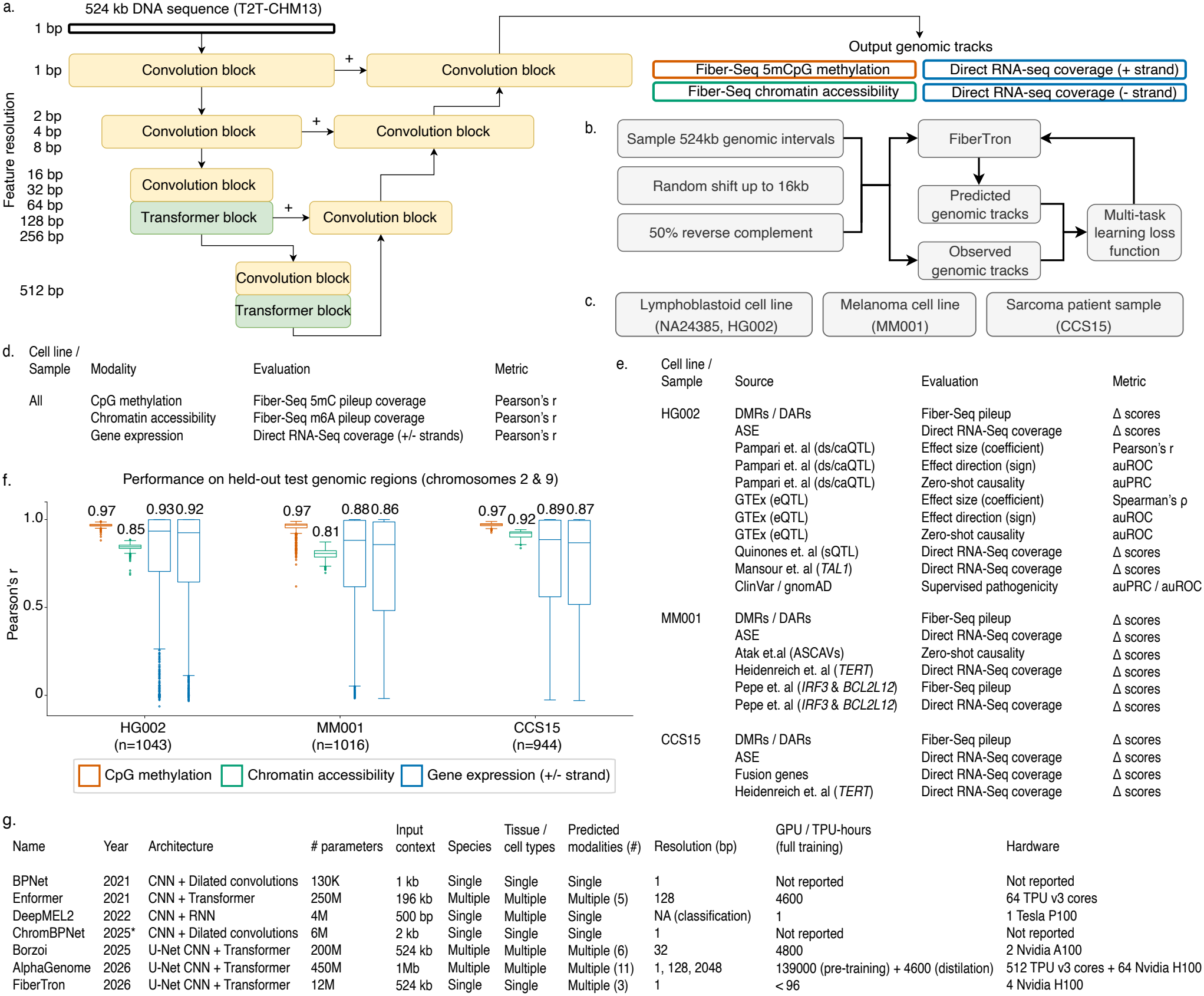
FiberTron: a lightweight multi-modal single-class sequence-to-function deep learning architecture for prediction and interpretation of Fiber-seq and direct RNA-seq datasets. **a.**Simplified schematic representation of FiberTron’s architecture (convolutional U-net backbone^9^ with hierarchical self-attention modules). **b**. Adopted supervised training framework; 524 kb DNA genomic intervals are sampled from the training set (T2T-CHM13v2.0 reference genome), augmented (shifted and/or reverse complemented) and used to train FiberTron instances against experimental targets (Fiber-seq and direct RNA-seq modalities). **c**. Cell lines/samples used for FiberTron instances’ training. **d**. Experimental track predictions. **e**. Outline of variant effect predictions. **f**. Box plots showing FiberTron’s genome-wide performance per modality across all genomic intervals in held-out test chromosomes (chr2 & chr9) for every cell line/sample used for training. Boxplots show the median (center line) and interquartile range (IQR; 25^*th*^–75^*th*^ percentiles). Whiskers extend to the most extreme values within 1.5*×*IQR of Q1 and Q3; points beyond are plotted as outliers. **g**. Table comparing FiberTron implementation to existing state-of-the-art (SoTA) models (*: bioRxiv preprint).

For the current study, three FiberTron instances (*≈* 12M parameters per instance) were trained in a supervised setting (Figure 1b) to decipher the genomic regulatory code of a human lymphoblastoid cell line (HG002, GM24385/NA24385) from the Genome-in-a-Bottle (GIAB) consortium^27^, a human melanoma cell line (MM001) and a patient-derived sample from a rare clear cell sarcoma (CCS15) (Figure 1c). Each FiberTron model instance was trained on experimental target vectors obtained from two sequencing experiments; a Fiber-seq run generated at *≈* 30*×* coverage and a single direct RNA-seq flow cell (*≈* 20M reads). The performance of FiberTron instances was evaluated at various track and variant effect predictions (Figure 1d-e) at held-out genomic regions, in-sample candidate *cis*-regulatory variants, and variant effect data from independent sources that were not utilized in any step of the model training procedure. FiberTron is trained on CHM13v2.0 reference sequence only and never sees alternate alleles. In-sample heterozygous variants therefore provide a conservative test of its variant-effect predictions. Collectively, FiberTron’s lightweight architecture and multi-modal training paradigm establish a departure from existing SoTA models, enabling accurate and scalable regulatory sequence interpretation while reducing computational complexity (Figure 1g).

Chromosome-wide statistics on held-out chromosomes demonstrate that FiberTron accurately predicts DNA methylation, chromatin accessibility and gene expression. Across all samples, predictions match the ground truth data with median Pearson’s *r* of 0.97, 0.85, 0.89 and 0.89 for methylation, accessibility and plus/minus-strand expression, respectively (Figure 1f). To complement the genome-wide statistical measures, the following subsections focus on specific genomic loci for which the characteristic patterns of the Fiber-seq and direct RNA-seq modalities are well established (e.g., CpG islands, CTCF binding sites, transcription start sites, promoters, enhancers, etc.). Unless explicitly indicated, the performance metrics reported in the subsequent sections are derived from held-out data from the HG002 cell line.

### FiberTron accurately predicts CpG methylation states

In the human genome, CpG methylation is an important epigenomic modification involving the addition of a methyl group to the 5^*th*^ carbon of cytosines followed by guanines to form 5-methylcytosines (5mCpG). Promoters and enhancers frequently overlap CpG islands. The methylation state of these CpG islands correlates with gene expression (methylated islands are typically associated with repression^28^) and changes are common in disease states, particularly cancer^29,30^. On the held-out chromosomes, FiberTron predicts both island-level and single CpG methylation rates with mean Pearson’s *r* correlation values of 0.861 (Figure 2a) and 0.789 (Figure 2b) respectively. Binarizing the methylation rates, FiberTron correctly classifies 91.23% and 70.01% of all methylated and unmethylated CpG’s, respectively (Figure 2c), achieving an accuracy of 0.847, precision of 0.872, recall of 0.912, and F1-score of 0.891 (Figure 2d). Across CpG islands, the predicted 5mC methylation profiles show high concordance with the ground truth with mean Pearson’s *r* correlation of 0.896 (Figure 2e).

**Figure 2.**
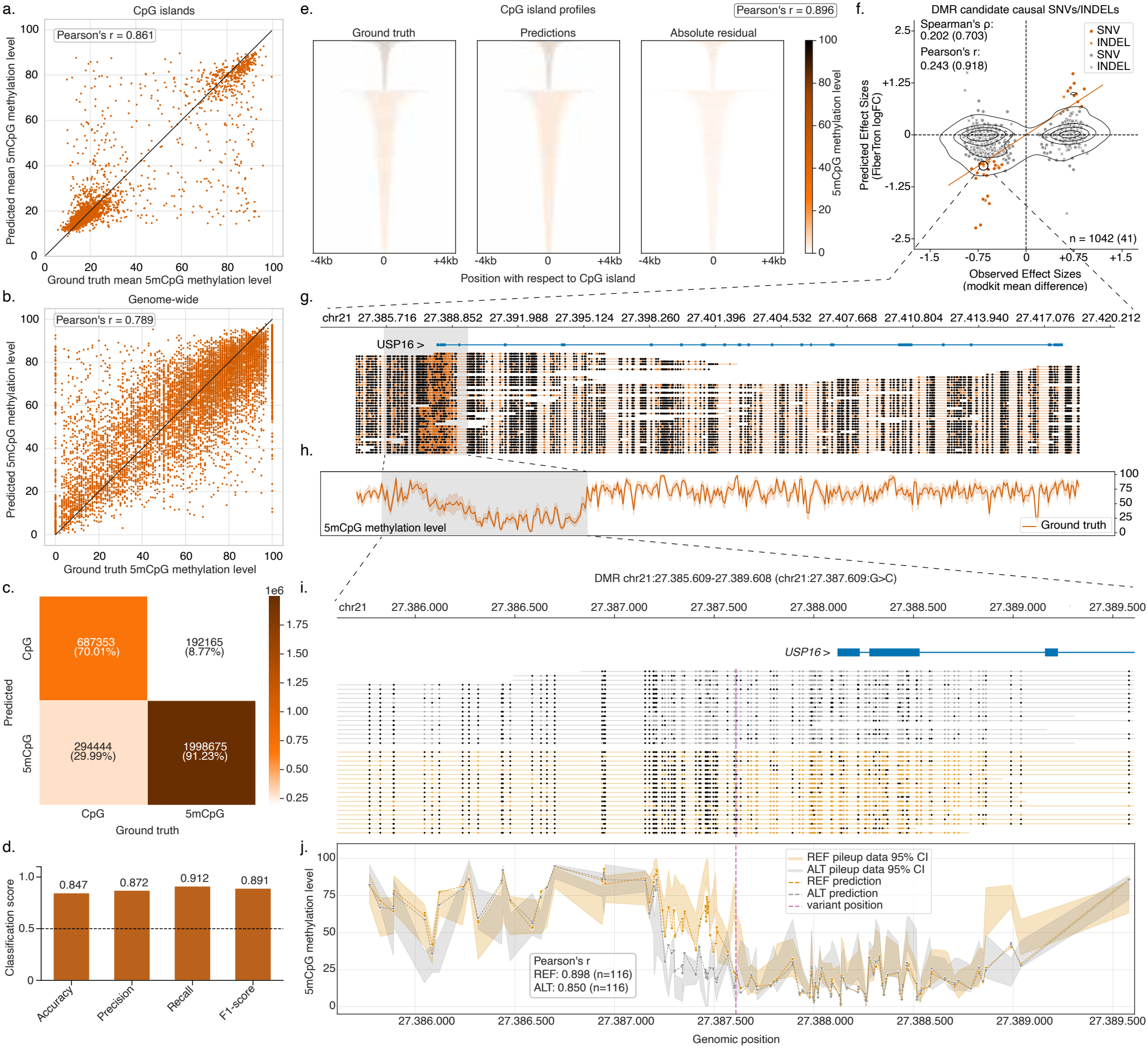
FiberTron predicts (allele-specific) CpG methylation from sequence. **a.**Comparison between predicted and ground truth mean 5mC methylation levels across all CpG islands in held-out chromosomes (Pearson’s *r* = 0.861). **b**. Comparison between predicted and ground truth genome-wide CpG methylation levels (10k randomly sampled CpG dinucleotides from held-out chromosomes, Pearson’s *r* = 0.789). **c-d**. Methylated/unmethylated binary classification task (**c**) confusion matrix and (**d**) performance metrics for all held-out CpG dinucleotides genome-wide. **e**. Ground truth, predicted, and absolute residual heatmaps of pileup 5mC methylation profiles at held-out CpG islands, split by binarized mean methylation level (threshold = 50) and sorted in descending order according to genomic span (mean Pearson’s *r* = 0.896). **f**. Scatter plot of observed allele-specific differentially methylated region (DMR) effects and FiberTron-predicted variant effect scores. Putatively causal *cis*-acting variants are depicted in orange color. **g**. Methylartist^33^ depiction of single-molecule PacBio HiFi reads aligned to the *USP16* gene locus. Methylated and unmethylated CpGs are shown as black and orange dots, respectively. The 4 kb region centered around the identified DMR is shaded. **h**. Mean 5mC methylation level curve (and 95% confidence interval bands) at the *USP16* gene locus. **i**. Same as (**g**), but zoomed in to the *USP16* promoter region and with reads colored by their assigned haplotype (yellow for REF or gray for ALT). Methylated CpGs are colored black. The purple vertical dashed line shows the chr21:27.387.609:G>C variant position. **j**. Allele-specific predicted 5mC methylation profiles (yellow dashed line for REF and gray dashed line for ALT) overlaid with the 95% confidence interval of the allele-specific pileup data. The purple vertical dashed line shows the chr21:27.387.609:G>C variant position. For **a-e**, all visualizations and reported performance metrics are based on held-out HG002 data.

*Cis*-regulatory variation may lead to allele-specific changes in CpG island methylation and consequent alterations in gene expression^31^. To test whether FiberTron can capture the effects of such regulatory variation, we evaluated its performance at in-sample allele-specific differentially methylated regions (DMRs). Using the phased long-read Fiber-seq reads, modkit^32^ identified 2385 DMRs in the HG002 sample, 1042 of which contained phased heterozygous variants, resulting in 4481 unique DMR-variant pairs. By using FiberTron to compute a variant effect score for each DMR–variant pair, we identify putatively causal *cis*-acting variants (Figure 2f). The *USP16* promoter contains a representative DMR including chr21:27387609:G>C variant (Figure 2g-j). Examination of the raw allele-specific methylation profiles (Figure 2i) revealed a *cis*-acting effect associated with the REF (G) and ALT (C) haplotypes. Across this locus, FiberTron’s allele-specific predictions showed strong agreement with both REF- and ALT-containing pileup data, achieving Pearson’s *r* values of 0.898 and 0.850 respectively (Figure 2j) and accurately recapitulated the direction and magnitude of the methylation difference between the two haplotypes. Specifically, FiberTron predicted reduced methylation levels for the ALT allele, in agreement with the observed methylation pattern and a trend to increased *USP16* expression for ALT relative to REF allele (121 versus 98 raw allele-specific counts; one-sided binomial test, p-value = 0.0685). These results illustrate that FiberTron can accurately recapitulate biologically meaningful allele-specific methylation differences associated with gene regulatory variation.

### FiberTron recapitulates base-pair level chromatin accessibility and long-range effects

To assess in detail the ability of FiberTron models to distill functional, specific chromatin accessibility patterns, we first focused on loci displaying characteristic nucleosome arrangements, such as CTCF binding sites and TSSs. Performance was evaluated within a 2 kb window around held-out CTCF binding sites and TSSs. FiberTron’s base-pair level chromatin accessibility predictions match ground truth Fiber-seq 6mA pileup profile data with mean Pearson’s *r* values of 0.842 and 0.813 at CTCF sites and TSSs, respectively (Figure 3a-b). At CTCF sites, the model accurately predicts the stereotypic nucleosomal locations, with two central peaks of accessibility enclosing the CTCF motif and footprint, reflected by a central focal dip in the accessibility. Likewise, predicted chromatin accessibility patterns at TSSs closely mirror those observed, and reflect the expression level category of the corresponding gene, suggesting that FiberTron distinguishes active from inactive genes (Figure 3b, Extended Data Figure 2a-d).

**Figure 3.**
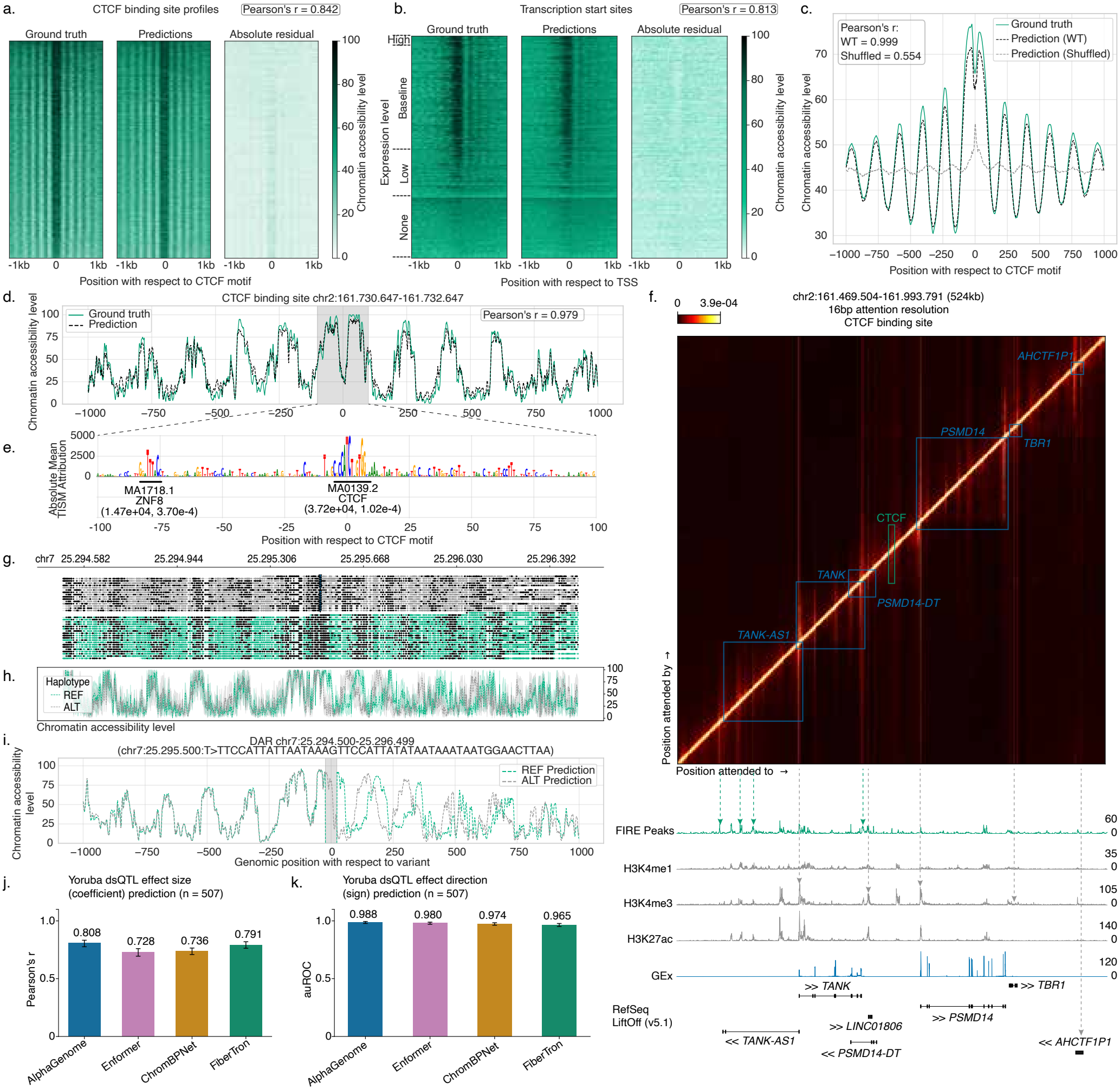
Distilling base-pair level chromatin accessibility and organization. **a-b.**Fiber-seq ground-truth, predicted, and absolute residual heatmaps of chromatin accessibility in 2 kb windows centered on (**a**) CTCF binding sites (ordered by decreasing mean signal intensity, mean Pearson’s *r* = 0.842) and (**b**) TSSs (ordered by decreasing gene expression count and mean signal intensity, mean Pearson’s *r* = 0.813). **c**. Mean chromatin accessibility profiles in 2 kb windows around CTCF binding sites for Fiber-seq ground truth and FiberTron predictions on wild-type and CTCF motif-shuffled sequences. **d**. Comparison between ground truth pileup and predicted chromatin accessibility profiles in a 2 kb window around a CTCF binding site at chr2:161.731.637-161.731.657 (Pearson’s *r* = 0.979). **e**. TISM^34^ ACWM for 200 bp sequence containing the example CTCF binding site. Tangermeme-identified^36^ seqlets are annotated (attribution score and p-value in parentheses). **f**. Heatmap visualization of the attention weight matrix for the 524 kb window centered on the example CTCF site. Annotation tracks showing FIRE elements, ENCODE^39^ ChIP-seq (GM12878) and gene expression (GEx) are plotted below. **g-i**. Aligned views of the chr7:25.294.500-25.296.499 region containing an allele-specific differentially accessible region (DAR) adjacent to a CTCF site and a 47 bp insertion (centered). (**g**) Methylartist^33^ depiction of Fiber-seq reads phased to the REF (green) or ALT (grey) alleles. The insertion is visible as a column of blue triangles on the top (ALT) haplotype. 6mA bases are shown as black dots. (**h**) Allele-specific pileup of the Fiber-seq 6mA chromatin accessibility signal at the site. (**i**) Predicted chromatin accessibility profiles for REF and ALT (insertion-containing) alleles. The position of the indel is highlighted. **j-k**. Comparative evaluation of FiberTron for predicting the effect size (**j**) and directionality (**k**) of Yoruba ancestry dsQTLs^40^. For **a-f**, except for ChIP-seq tracks, all visualizations and reported performance metrics are based on held-out HG002 data.

Binding of CTCF to its cognate motif on DNA is necessary for the local organization of nucleosomes. To test whether FiberTron functionally understands the importance of CTCF binding motifs, we randomly shuffled them (15 bp) and assessed model predictions. After shuffling, predicted local chromatin accessibility no longer displayed the typical CTCF-induced patterns (Figure 3c), confirming that FiberTron has internalized the impact of these motifs within their genomic contexts. Even at individual CTCF binding sites, FiberTron accurately charts chromatin accessibility profiles at base resolution and offers detailed insights into their regulatory grammar (Figure 3d; example was selected based on the overall shape of the ground truth pileup to obtain a clear CTCF-induced pattern and on the concordance between the ground-truth and predicted 6mA profiles). Using a gradient-based approximation for in silico saturation mutagenesis (ISM)^34^, an attribution-based contribution-weight matrix (ACWM)^35^ can be derived for the entire 524 kb input sequence centered around the example CTCF binding site. Tangermeme seqlet calling^36^ on a ACWM slice corresponding to 2 kb around the CTCF binding site recovers the known CTCF binding motif (JASPAR^37^ entry MA0139.2; Figure 3e).

CTCF is known to have long-range chromatin organizational effects, and the example binding site is located at the center of a gene-rich locus with 7 genes arrayed across the 524 kb genomic region around it (chr2:161.469.504-161.993.791). To better understand which parts of the input sequence FiberTron focuses on to make predictions for the whole locus the attention weight matrix can be explored (e.g., at 16 bp resolution; Figure 3f). The presence of strong off-diagonal attention signals demonstrates that FiberTron attends to distant sequences for its prediction. Moreover, the loci attended to closely correspond to regulatory elements as evidenced by peaks of Fiber-seq inferred regulatory elements (FIRE)^38^ (HG002) as well as orthogonal ENCODE^39^ tracks of H3K4-methylation and H3K27 acetylation (GM12878), but also genic elements (e.g., exons) and the central CTCF site. Upon shuffling the CTCF motif, the pronounced local off-diagonal signal is no longer observed in FiberTron’s attention map, indicating the site is no longer attended to (Extended Data Figure 2e). Furthermore, the CTCF binding motif is no longer highlighted in FiberTron’s ACWM (Extended Data Figure 2f). These findings demonstrate that FiberTron captures and integrates biologically relevant sequence determinants of chromatin accessibility and organization, both locally and hundreds of kb away.

Next we sought to investigate whether FiberTron can predict the impact of genetic variation on chromatin accessibility. As FiberTron is trained on reference genome sequence alone, we can directly test and validate predictions using in-sample differentially accessible regions (DARs) and (candidate causal) *cis*-regulatory variants identified from the Fiber-seq data. Analogous to the DMR analysis, modkit^32^ identified 923 DARs, of which 371 harbored phased heterozygous variants, yielding 1665 unique DAR–variant pairs. One such DAR was identified near a CTCF binding site at chr7:25.294.500-25.296.499 and contains a 47 bp insertion (ALT) (Extended Data Figure 2g). Exploring the phased Fiber-seq reads at the locus reveals how the DAR may arise due to interactions between the local chromatin structuring imposed by CTCF and the 47 bp sequence offset introduced by the indel. Specifically, nucleosome positions (dips in the chromatin accessibility) are shifted by approximately 47 bp downstream of the CTCF site, but remain unaltered upstream (Figure 3g-h). FiberTron’s predictions for the REF and ALT alleles indicate that the model accurately captures these effects (Figure 3i). Further illustrating FiberTron’s predictive capabilities, relocating the indel in silico upstream of the CTCF binding site and separately replacing it with a random insertion (lacking a CTCF binding motif) likewise shows the corresponding expected shifts in nucleosome positioning (Extended Data Figure 2h-i). Note that, while these interactions may be expected, they are non-obvious, and likely to be missed using short-read approaches (lack of phasing) and models trained on such data. In contrast, FiberTron has captured the genomic syntax dictating short and longer range chromatin organization.

Additionally, to evaluate the model’s capacity to capture the functional consequences of genetic variation, we employed two external QTL datasets: the Yoruba ancestry dsQTL^40^ and the African ancestry caQTL^41^ sets. We assessed FiberTron’s performance in predicting both the magnitude (coefficient) and direction (sign) of the reported effect sizes, as well as its ability to discriminate putatively causal variants from non-causal ones in a zero-shot mode. FiberTron predicts the observed effect sizes with Pearson’s *r* values of 0.791 and 0.638 and directions with auROC values of 0.965 and 0.871 (Extended Data Figure 3a, c) for the dsQTL and caQTL sets, respectively. Making use of FiberTron’s multi-modality, predicted effects for CpG methylation anti-correlate with chromatin accessibility, as expected (Extended Data Figure 3b, d).

Trained on a single HG002 Fiber-seq and direct RNA-seq data set as opposed to large-scale consortium data, FiberTron recovers dsQTL effect magnitudes and directions to within 0.017 Pearson’s *r* and 0.023 auROC of the best ChromBP-Net^5^, Enformer^8^ and AlphaGenome^10^ models trained on cell type-matched DNase- and ATAC-seq (Figure 3j-k, Extended DataFigure 3e-f), but is outperformed on the African-ancestry caQTL set (Pearson’s *r* = 0.638 versus 0.733, auROC = 0.871 versus 0.952; Extended Data Figure 3h-i). It is important to note that comparator models predict the chromatin profiling assays (DNase-/ATAC-seq) against which they are scored, whereas FiberTron is trained on materially distinct Fiber-seq 6mA footprinting data, which is likely to affect these numbers. When it comes to identifying putatively causal variants, FiberTron’s performance is on par with ChromBPNet models trained on ATAC-seq data downsampled to 25M reads (Extended Data Figure 3g).

To further assess the extent to which the individual FiberTron instances learn biologically meaningful features and internalize the underlying regulatory landscape of the respective samples, we evaluated model performance at loci exhibiting cell-type-specific chromatin accessibility. In contrast to previous reports showing diminished performance of SoTA foundational models in cell-type-specific accessible regions^42^, FiberTron, as expected due to its single-sample training framework, did not display a strong systematic performance bias associated with such regions across any of the cell lines on which individual model instances were trained (Extended Data Figure 4). Across all examined genomic region spans centered on DNaseI/FIRE peaks, FiberTron predictions for chromatin accessibility exhibited consistent performance across cell-type specificity bins, as quantified by the mean Pearson’s *r* (Extended Data Figure 4a-d). For 501 bp genomic regions, prediction performance across cell-type specificity bins showed only weak ordinal associations with decreasing cell-type specificity (Kendall’s *τ* ∈ [*−*0.04, 0.16], Extended Data Figure 4e-h), indicating that FiberTron’s accuracy remained largely stable across the different levels of cell-type specificity.

### FiberTron accurately predicts gene expression and learns relevant motifs

Predicting gene expression from genomic sequence alone poses distinct challenges. First, (direct) RNA-seq coverage reflects the result of multiple processes, including transcription, splicing, polyadenylation and degradation. Second, in contrast to the mostly local effects of CpG methylation and chromatin accessibility, gene expression often involves interactions between regulatory elements that can be several kilobases or even megabases apart. To assess the ability of FiberTron to accurately model the intricacies of gene expression we focus on protein-coding genes whose entire length is included in FiberTron’s output range (328 kb, 1735/2035 genes on held-out chromosomes or 85.3%; Figure 4a). Stratified by gene expression level, FiberTron achieves median Pearson’s *r* values of 0.99, 0.69, 0.91 and 0.95, respectively, for silent, lowly, mid and highly expressed genes. In contrast, performance for CpG methylation and chromatin accessibility is similar across all categories (median Pearson’s *r* values of 0.96 and 0.84, respectively). At the level of transcript counts, FiberTron log_2_-normalized raw count predictions (+1 pseudocount) match the HG002 direct RNA-seq ground truth with a mean Pearson’s *r* value of 0.815 across its broad dynamic range (Figure 4b).

**Figure 4.**
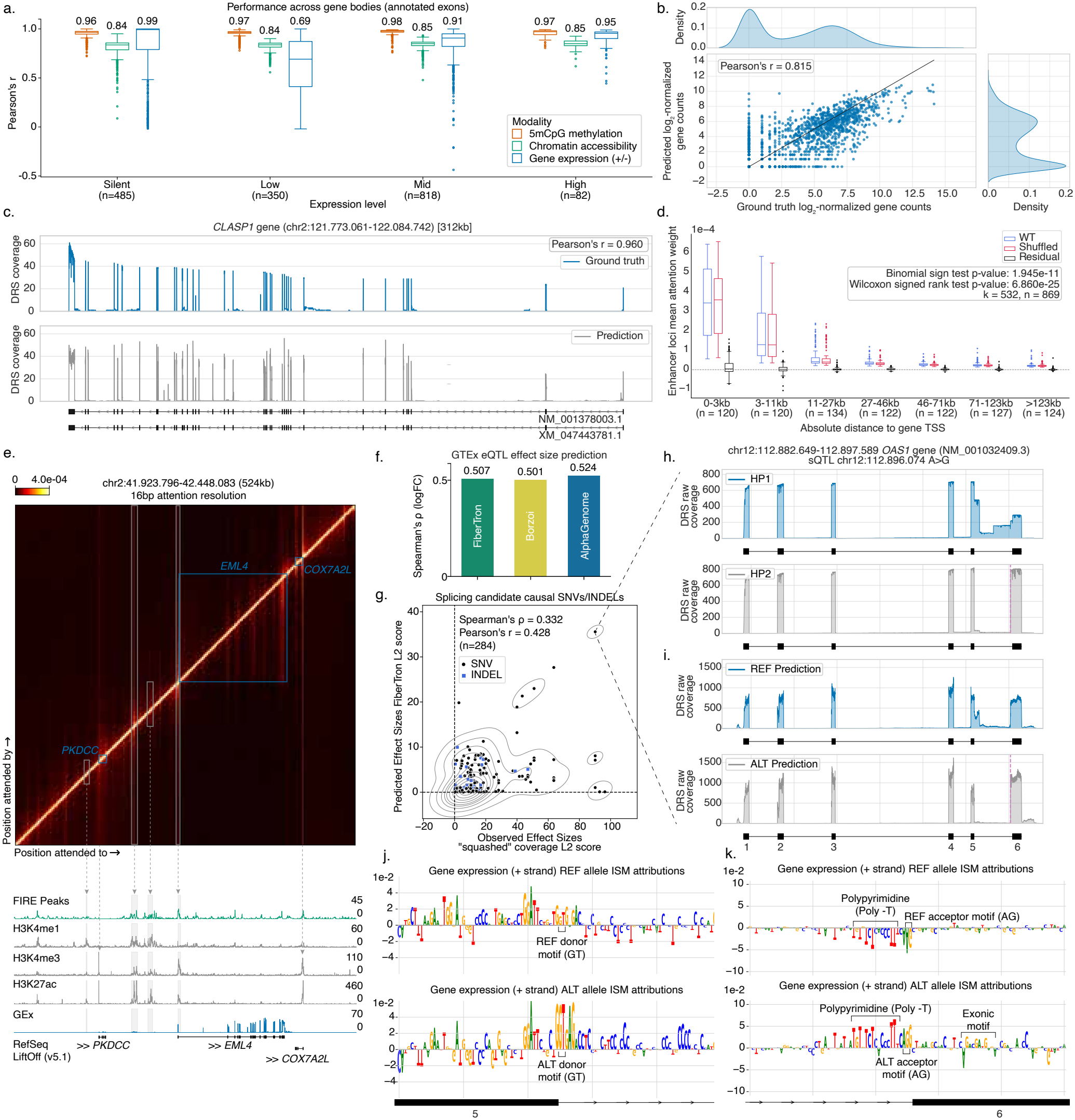
Predicting gene expression and deciphering transcriptional syntax. **a.**Correlation between FiberTron predictions and Fiber-seq/direct RNA-seq ground truth across gene bodies on held-out chromosomes, stratified by expression level. **b**. Comparison of direct RNA-seq observed and FiberTron-predicted log_2_-normalized gene counts. **c**. Observed and predicted gene expression coverage profiles for *CLASP1*. **d**. Box plots of mean attention weight given by wild-type and shuffled gene promoters to enhancer loci stratified by distance from the TSS. Residual attention (WT - Shuffled) is shown in black. **e**. Attention weight matrix for the *EML4* locus. FiberTron attends to ENCODE-annotated enhancers interacting with the *EML4* promoter. **f**. GTEx eQTL^44^ effect size prediction performance of FiberTron, Borzoi^9^, and AlphaGenome^10^. **g**. Comparison of FiberTron-derived and direct RNA-seq observed L2 scores for the prioritization of putative causal variants driving allele-specific splicing. **h-i**. (**h**) Ground truth direct RNA-seq and (**i**) FiberTron-predicted *OAS1* allele-specific transcript coverage. The causal chr12:112.896.074:A>G (rs10774671) variant is indicated by a dashed vertical line. **j-k**. ISM ACWMs for REF and ALT allele gene expression for (**j**) a 101 bp window centered on the exon 5 donor splice site and (**k**) a 101 bp window centered on the splicing variant at exon 6. For **a-e**, except for ChIP-seq tracks, all visualizations and reported performance metrics are based on held-out HG002 data.

While FiberTron is never provided with gene annotations, it accurately recapitulates direct RNA-seq transcript coverage patterns across entire genes, including exon-intron patterns, even for long genes (e.g., 312 kb *CLASP1*; Figure 4c). To achieve this, FiberTron is expected to integrate long-range regulatory dependencies between distal *cis*-regulatory elements (cREs) and their cognate target genes. To test this, we systematically evaluated attention allocation at loci corresponding to annotated enhancer-promoter interaction pairs in FiberTron attention maps and compared these patterns to those obtained after promoter sequence shuffling (Methods). For 532 out of 869 putative interaction pairs located on held-out chromosomes, FiberTron assigned higher attention scores to the wild-type sequence compared to their shuffled counterparts (binomial sign test, p-value = 1.945 *×* 10^*−*11^) (Figure 4d). This is likely an underestimate, as high-throughput chromosome conformation capture (Hi-C) assays^43^ primarily identify physical contacts between enhancers and promoters, which do not necessarily correspond to functional regulatory interactions influencing transcriptional output. In contrast, in line with its training objective, FiberTron may therefore prioritize sequences directly impacting gene expression. Indeed, attention weight matrices reveal how FiberTron focuses on ENCODE-annotated cREs^39^ (ChIP-seq tracks for lymphoblastoid line GM12878) when predicting functional modalities across the representative *EML4* gene locus (Figure 4e). Notably, these cREs overlap biologically relevant *EML4* enhancers located approximately -141 kb, -71 kb, and -46 kb from its TSS, further confirming that FiberTron leverages distal regulatory information to inform locus-specific predictions.

We next evaluate whether FiberTron can capture the influence of genetic variation on gene expression. To this end, we leveraged fine-mapped (i.e., putatively causal) expression quantitative trait loci (eQTLs) from the Genotype-Tissue Expression (GTEx) consortium (re-processed as per Wang *et al*.^44^). Across multiple evaluation tasks, FiberTron performs on par with substantially larger SoTA models, including Borzoi and AlphaGenome (Figure 4f, Extended Data Figure 5a-f). For eQTL effect-size prediction (SuSiE *β* posterior estimate), FiberTron achieves a Spearman’s *ρ* of 0.507, compared with 0.501 for Borzoi and 0.524 for AlphaGenome (Figure 4f, Extended Data Figure 5b). Furthermore, FiberTron infers eQTL effect directionality (sign of the SuSiE *β* posterior estimate), attaining an auROC of 0.779 versus 0.797 and 0.811 for Borzoi and AlphaGenome, respectively (Extended Data Figure 5c-d). Finally, in a zero-shot mode, FiberTron-derived variant effect scores (Methods) effectively distinguish putatively causal variants from randomly sampled variants. FiberTron achieves an auROC of 0.718 compared with 0.781 and 0.809 for Borzoi and AlphaGenome, respectively (Extended Data Figure 5e-f), indicating that FiberTron scores inform on regulatory variant function. Collectively, these results demonstrate that FiberTron captures both the magnitude and direction of regulatory variant effects on gene expression and enables effective prioritization of putatively causal variants.

Further probing FiberTron’s understanding of gene expression, we also assess in-sample allele-specific splicing. We first detect allele-specific exon usage in our HG002 direct RNA-seq data and subselect to genes with *cis*-directed alternative splicing events reported in the related GM12878 lymphoblastoid cell line^45^. Using an L2-based score, FiberTron-predicted versus direct RNA-seq observed *cis*-directed splicing effect sizes showed a Spearman’s *ρ* of 0.332 (Figure 4g). Interestingly, the top hit, chr12:112.896.074:A>G (rs10774671) is a well-known COVID-19 risk variant in *OAS1*. Disruption of an efficient exon 6 splice acceptor site, giving rise to the protective p46 isoform, by the A allele yielding multiple alternative isoforms p42, p44a/b, p48 and p52 (Figure 4h, Extended Data Figure 6a)^46,47^. Despite the complexity of these coordinated changes, they are remarkably well recapitulated by FiberTron’s allele-specific predictions (Figure 4i, Extended Data Figure 6a).

As before, FiberTron models allow for biological interpretation and data exploration. Specifically, in silico saturation mutagenesis (ISM) using FiberTron around the two splice junctions involved can provide mechanistic insights into the local regulatory syntax and the effects of the *OAS1* splice variant. The TCA[A>G]GCT variant creates a strong splice acceptor, directing effective splicing, as highlighted by the positive attribution scores for the acceptor motif and the adjacent polypyrimidine and exonic motifs (Figure 4j-k). In contrast, transcripts from the A allele utilize various cryptic splice donor and acceptor sites up- and downstream of the variant, encoding a wide array of alternate isoforms (Extended Data Figure 6a). These patterns are similarly visible in the ACWMs (Figure 4j-k, Extended Data Figure 6c). Presence of the strong acceptor similarly activates splicing at the canonical upstream 5’ donor site, reflected in increased attribution scores observed in the ISM for the alternative allele (Figure 4j bottom panel) compared to those for the reference allele (Figure 4j top panel). Taken together, these findings show that FiberTron can accurately recapitulate intricacies of gene expression on human chromatin and may offer mechanistic insights into the effects of regulatory variation, including those affecting transcriptional processes.

### FiberTron captures the effects of small, complex and disease-associated variants

To evaluate the biological and clinical relevance of FiberTron variant effect scores, we assessed whether the HG002 model can discriminate benign from pathogenic variants. To this end, we use the ClinVar^48^ database annotations and common (likely benign) versus rare (potentially deleterious) variants in the gnomAD^49^ database. FiberTron-derived effect scores achieved a mean 10-fold cross-validated auPRC of 0.739 across all ClinVar molecular consequence categories (Extended Data Figure 7a), corresponding to a 2.5-fold improvement over a random baseline (Extended Data Figure 7b). For intronic and synonymous variants, we observed a 20-fold increase in performance. For gnomAD rare variants, FiberTron-derived scores outperformed CADD (v1.7)^50^ RAW and PHRED scores, achieving a mean 10-fold cross-validated auROC of 0.706 compared to 0.625 (Extended Data Figure 7c). Including CADD scores further improves predictive performance (auROC = 0.726), likely reflecting the inclusion of protein-altering variant effect knowledge complementing FiberTron’s gene regulatory insights.

We next evaluated whether FiberTron can capture in-*cis* effects of genetic variation in clinically relevant disease states. To this end, we trained FiberTron model instances on matched Fiber-seq and direct RNA-seq data from the patient-derived melanoma cell line MM001 and fresh-frozen tissue from a rare case of clear cell sarcoma, CCS15. Utilizing a previously reported ground truth set^6^, we tested whether FiberTron chromatin accessibility effect scores can reliably distinguish causal allele-specific chromatin accessibility variants (ASCAVs) from random variants in the melanoma line MM001. FiberTron-derived scores explain 79% of the observed ASCAVs (n=1,840) at a 0.05 false positive rate (Figure 5a). Of these, 56.4% (44.6% of total) could additionally be mechanistically linked to changes in known TF binding motifs via FiberTron attribution TISM and seqlet mapping (Extended Data Figure 8a). This represents a substantial increase in variant explainability relative to the *≈* 10% in the original DeepMEL2 study reporting the benchmark^6^. Note that, in contrast to FiberTron’s single-sample training framework, in DeepMEL2 a limited-context convolutional recurrent DL architecture was trained on ATAC-seq profiles from 30 melanoma samples. The comparative analysis of accessible versus inaccessible alleles highlighted biologically meaningful motifs affected by ASCAVs, including ETS-, SOX- and MYB-family transcription factors (enriched in other melanocytic proliferative melanomas^6^). While these motifs were not enriched in accessible or inaccessible alleles (Extended Data Figure 8b), they were overrepresented in ASCAVs, among other biologically relevant motifs, relative to control variants (Extended Data Figure 8c).

**Figure 5.**
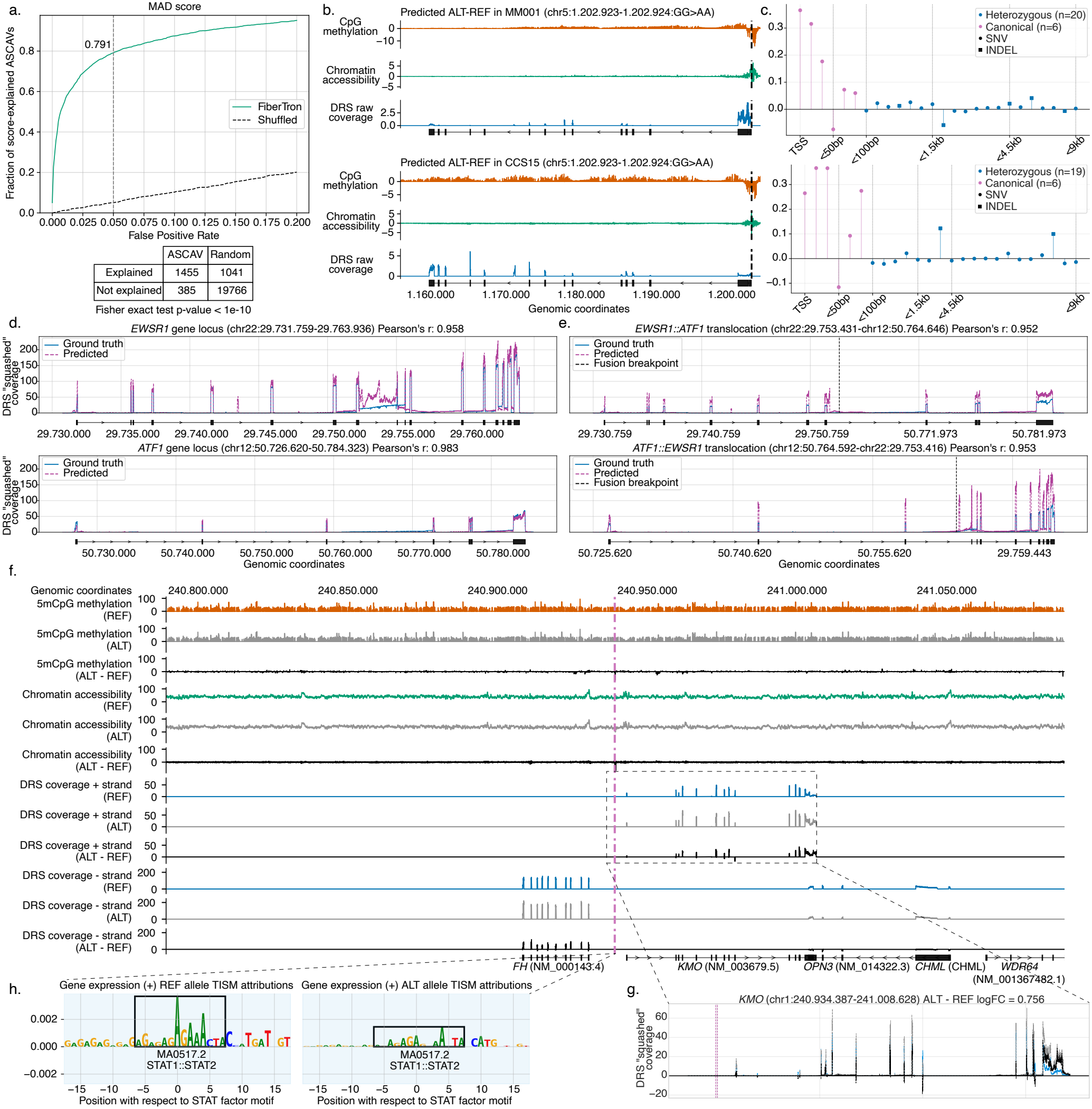
Modeling effects of complex and disease-associated genetic variants. **a.**Fraction of score-explained allele-specific chromatin accessibility variants (ASCAVs) in melanoma line MM001 at different false positive rates (FPR). The 2 *×* 2 contingency table for FPR=0.05 is shown. **b**. FiberTron-predicted Δ effects (ALT - REF) of a canonical *TERT* promoter mutation in melanoma line MM001 line (top) and clear cell sarcoma case CCS15 (bottom). **c**. Predicted effect scores for canonical driver and heterozygous in-sample variants in the wider *TERT* promoter region for MM001 (top) and CCS15 (bottom). **d-e**. Comparison of direct RNA-seq ground truth and FiberTron-predicted gene expression coverage for (**d**) *EWSR1, ATF1* and (**e**) their reciprocal fusion genes *EWSR1::ATF1* and *ATF1::EWSR1* in clear cell sarcoma sample CCS15. **f**. Reference, alternate and Δ (ALT - REF) predicted profiles for a *≈* 328kb window around the *FH* and *KMO* locus, containing a candidate causal 334 bp deletion at chr1:240.939.801-240.940.135 in HG002 cell line. **g**. Magnified view of the predicted *KMO* gene expression coverage on the reference and alternative alleles and their difference. **h**. TISM^34^ ACWMs around the variant for the reference and alternate allele shows differential attribution score magnitudes for STAT-family transcription factor (TF) motifs.

Non-coding cancer driver mutations are notoriously hard to identify, and as a result, few have been described^51^. Our FiberTron framework may aid their characterization, as both training and initial validation data as well as mechanistic insights may be derived from a single clinical sample profiled with Fiber-seq and direct RNA-seq. Mutations activating the *TERT* promoter are some of the best-characterized non-coding drivers and are found in various cancer types^52^, including melanoma^53^. Interestingly, clear cell sarcoma CCS15 carries a non-canonical *TERT* promoter mutation (chr5:1207403:G>A)^54^ with unknown function. To determine its functional relevance as a passenger versus likely driver mutation, we derive MM001 and CCS15 FiberTron effect scores for all in-sample heterozygous variants within 10 kb of the *TERT* promoter and include the canonical promoter mutations. FiberTron indicates an increase in *TERT* gene expression and corresponding promoter accessibility as well as changes to CpG methylation (gene body hyper- and promoter hypomethylation) for 5 and 4 of the 6 canonical mutations in MM001 and CCS15 respectively, in line with literature evidence (Figure 5b-c)^52^. Low effect scores for in-sample variants suggest they act as passengers in the two cancer samples (Extended Data Figure 7d-e). In Figure 5b the effects of the *TERT* promoter mutation with the highest predicted Δ score (chr5:1.202.923-1.202.924:GG>AA) are depicted. For both MM001 (top panel) and CCS15 (bottom panel) the variant results in a decrease in methylation and increase in chromatin accessibility of the *TERT* promoter locus, an increase in methylation of the *TERT* gene body, and increase in its expression. ISM ACWMs for the strongest canonical driver chr5:1.202.923-1.202.924:GG>AA mechanistically implicate ETS-like binding motifs in *TERT* activation, in line with previous findings^52^ (Extended Data Figure 9a-b).

Beyond *TERT* promoter mutations, FiberTron accurately recapitulates the regulatory effects of 17 out of 20 *TAL1* neo-enhancer mutations^55,56^ (Extended Data Figure 7f), as well as 2 out of 3 previously reported functional mutations associated with altered methylation and gene expression at the *IRF3* and *BCL2L12* loci^57^ (Extended Data Figure 7g-h). CCS15 additionally contains a t(12;22)(q13;q12) balanced translocation generating the *EWSR1::ATF1* fusion gene characteristic of clear cell sarcoma^58^. To assess FiberTron performance at this unseen fusion gene, we used the loci encoding the intact *EWSR1, ATF1* and the reconstructed derivative *EWSR1::ATF1* (and its reciprocal *ATF1::EWSR1*) to re-align the CCS15 direct RNA-seq and predict gene expression coverage. FiberTron accurately predicts transcript coverage of both *EWSR1* and *ATF1* (Pearson’s *r* = 0.958 and 0.983 to ground truth direct RNA-seq, respectively; Figure 5d) but also recapitulates the intron-exon structure and expression level of the fusion and its reciprocal (Pearson’s *r* = 0.952 and 0.953; Figure 5e). The oncogenic activity of *EWSR1::ATF1* in clear cell sarcoma has been linked to its recruitment to distal regulatory elements enriched for TF binding signatures from the CREB, AP-2, and SOX families^59^. Consistent with this observation, gradient-based FiberTron ACWMs and seqlet analysis revealed enrichment of motifs belonging to these families in CCS15 compared with both MM001 and HG002 (Extended Data Figure 8d). To place these observations in a broader regulatory context, differential enrichment analysis was subsequently performed across all TF families detected in the samples (Extended Data Figure 8e-f). This analysis further supported the preferential representation of CREB, AP-2, and SOX family motifs in CCS15, with these families exhibiting positive enrichment relative to MM001 and HG002. For AP-2 and SOX family motifs, which are also characteristic of melanocytic (proliferative) melanoma cell lines such as MM001^6^, enrichment differences between CCS15 and MM001 were modest, with log_2_ fold-changes of 0.74 versus 0.64 and 0.77 versus 0.51, respectively. In contrast, comparisons with the normal diploid HG002 genome revealed stronger differences, with depletion of AP-2 and SOX family motifs at these loci (*−*2.05 and *−*1.82, respectively). Notably, CREB family motifs, which include the ATF1 fusion partner, exhibited the strongest enrichment in CCS15 (log_2_ fold-changes of 2.00, *−*1.13, and *−*1.53 in CCS15, MM001, and HG002, respectively).

Given its large context window, FiberTron allows functional profiling of structural variation (SV). To systematically assess FiberTron-derived scores for structural variants (Methods), we identified in-sample SVs with putative in-*cis* functional effects from the HG002 Fiber-seq data. One of the highest-scoring examples (Extended Data Figure 9c) is a 334 bp deletion at chr1:240.939.801–240.940.135, approximately 8.5 kb downstream of the *FH* gene and approximately 3 kb upstream of the *KMO* TSS. FiberTron predicts the deletion to increase expression of both genes (Figure 5f-g, Extended Data Figure 10c) which is supported by overexpression observed from the deletion haplotype in the direct RNA-seq data (*KMO*, Extended Data Figure 10a, 17 versus 36 raw allele-specific counts, one-sided binomial test p-value = 0.00274; *FH*, Extended Data Figure 10b, 328 versus 388 raw allele-specific counts, one-sided binomial test, p-value = 0.0137). Gradient-based attribution identifies multiple motifs belonging to the STAT family of transcription factors in close proximity to the SV (Figure 5h, Extended Data Figure 9d), as well as near the *KMO* promoter (Extended Data Figure 9e), in line with its observed regulation by JAK-STAT signaling and its role as an interferon-stimulated gene^60^.

## Discussion

We developed FiberTron, a 524kb-context S2F architecture for regulatory variant effect prediction and interpretation. FiberTron shows that a single sample profiled by Fiber-seq and direct RNA-seq provides enough information to train a large-context S2F model predicting genome-wide CpG methylation, chromatin accessibility, and gene expression at base pair resolution. With only *≈* 12M parameters, FiberTron instances can be trained from scratch in less than a day on a standard GPU node and perform competitively to SoTA models on 10 variant effect prediction tasks across modalities. As FiberTron models are trained on reference sequence only, the phased reads of the same sample remain available to test model predictions, while sequence attribution yields mechanistic insights by linking them to candidate modality-specific motifs.

Models such as Borzoi and AlphaGenome are trained on compendiums of public short-read tracks across cell types and species, now achieving base-resolution with up to a megabase of context. In contrast, lightweight models such as ChromBPNet are trained on one assay and offer only a few kilobases of input. FiberTron combines the long context and multiple modalities of the former with the single-sample training of the latter. Sample-specific models have been trained on short-read accessibility for personal genome interpretation^6,61^. Fiber-seq is used in MethylSeqNet^20^ and FiberFold^19^ to provide additional measured inputs (i.e., methylation, regulatory elements) to condition expression and accessibility outputs from a pretrained Borzoi model, and predict 3D genome organization, respectively. To our knowledge, FiberTron is the first sequence-only model trained end-to-end on long-read multi-omic data from a single sample, and the first to predict Fiber-seq accessibility, direct RNA-seq coverage and CpG methylation jointly at base resolution. FiberTron can score any variant fitting its 524 kb window, including large SVs and translocations as we show for the *EWSR1::ATF1* fusion. Trained on the telomere-to-telomere complete CHM13v2.0 reference with long-read targets, it could make predictions where short-read assays provide no data.

The HG002 FiberTron instance matches the performance of large compendium-trained SoTA models such as AlphaGenome in dsQTL (Yoruba) and eQTL (GTEx) prediction tasks. It fell behind on African-ancestry caQTLs and in zero-shot discrimination of causal chromatin accessibility variants. Differences between Fiber-seq 6mA footprints and short-read DNase- or ATAC-seq signals on which these QTLs are called (and the compendium models trained) may explain part of this.

FiberTron represents a key component of the recently coined “near-perfect genome sequencing (NPGS)”, where the convergence of long-read technologies, haplotype-resolved assembly with pangenome integration, and AI-driven variant interpretation hold clinical potential^62^. Indeed, while its long-read inputs simultaneously enable construction of near-complete personal genomes, FiberTron learns from the reference sequence and haplotype-averaged signals. Unlike models trained on allele-specific measurements^63^, this leaves the sample-specific genetic variation unseen and available to test the model’s variant effect predictions. As we demonstrate, this can effectively be done at heterozygous variants, by phasing the multi-omic signals from the input long reads, and is mostly infeasible with short reads. This in-sample validation facilitates variant prioritization but also has caveats: (1) we train on haplotype-averaged signal, so allelic differences are shrunk and predicted effects are conservative, (2) allelic differences can also arise from imprinting or somatic epimutations^38^, and (3) haplotypes differ at many linked variants, so allelic data alone cannot single out the causal one. By nominating the variant and motifs it disrupts, FiberTron models can pinpoint variant(s) that best explain haplotype differences. In-sample validation hence further prioritizes candidates for perturbation experiments rather than replacing them.

Single-sample training matters especially where consortium data are thin. Large multi-class S2F models demonstrate progressively lower predictive accuracy within increasingly cell-type-specific accessible chromatin regions, which are enriched for disease-associated variants^42^. While added capacity per cell type can partly recover from this bias, FiberTron instances, by construction, are in distribution for the sample they model^64^. As expected due to its single-sample training framework, FiberTron instances do not suffer from this bias and predictions show consistent performance across cell-type specificity bins. In line with this, FiberTron discriminates benign and pathogenic variation, and recapitulates tumor-specific regulation: 79% of all allele-specific accessibility variants in a melanoma line, but also oncogenic mutations in the *TERT* promoter, *TAL1* neo-enhancers, and the characteristic *EWSR1::ATF1* fusion in clear cell sarcoma.

Fiber-seq with full-length RNA sequencing remains an exception in routine diagnostics. Where it is generated, e.g., for a rare tumor, FiberTron yields a model of the sample and a test of its predictions. Elsewhere, instances trained on representative samples, such as the lymphoblastoid HG002, can score variants from routine sequencing. Precomputed scores for every possible single-nucleotide variant now exist^65^ to support clinical decision making, but only for cell types in the training compendium.

A number of limitations concern our training targets. First, FiberTron learns from piled-up signal rather than from individual fibers. While regulatory elements are actuated largely all-or-none on single fibers^14^, this status frequently differs between haplotypes within, and between, single HG002 cells^66^. A pileup therefore measures how often an element is actuated, but it does not resolve co-actuation of neighboring elements. This is further confounded when complex tissues containing multiple cell types are profiled. Second, Hia5 labels only adenines, so footprint resolution varies and any sequence biases in labeling or detection^14,67^ are likely learned alongside chromatin state, as uncorrected accessibility models learn Tn5 and DNaseI bias^5^. Third, non-genetic allelic imbalances, e.g., due to imprinting or somatic epimutations, cap the allelic recall of any S2F model, warranting additional inputs for their resolution^20^. Clonal expansions in cell lines and tumors increase the number of such detectable somatic epimutations. Conversely, imbalances where FiberTron predictions for the two haplotypes are neutral may nominate candidate epimutations. Fourth, commonly in cancer, dosage effects due to copy number alterations shape gene expression. S2F models however generally have no information on DNA copy number available, likely limiting their accuracy in copy number aberrated samples (note that only copy number-independent ASCAVs are used for evaluation in MM001). Fifth, FiberTron instances were trained on the data equivalents of a single flow cell each on a PacBio Revio (*≈* 30*×* whole-genome Fiber-seq coverage) and an ONT PromethION (*≈* 20M RNA reads), putting also generation of the required training data within reach of individual labs. Nevertheless, given the stochasticity of chromatin at the single-molecule level, the complexity and dynamic range of the transcriptome (e.g., prediction accuracy is reduced for lowly expressed genes), and the heterogeneity of tissues, we anticipate that increasing data may further improve model accuracy^5,20^.

Further challenges relate to the evaluation and scope of S2F models generally. Sequence homology can result in information leakage between training and held-out chromosomes, potentially inflating performance, and accuracy on reference sequence can overstate robustness to sequence edits^68,69^. Modeling the influence of distal (i.e., >100 kb) regulatory elements remains an ongoing challenge^10^. Our evaluations focused on protein-coding genes and HG002, with annotations transferred from the well-annotated GM12878 line, and include in-sample validation but no perturbation experiments. Although per-sample training is practical, the memory footprint of attention requires data-center GPUs (*<* 24 h on four H100 GPUs).

Future work may address several of these limitations. First, phased reads could supply allele-resolved targets to FiberTron in a contrastive learning framework^63,70^. Chromosomes would then need to be held out to preserve an in-sample test. Second, generative models predicting the state of individual fibers, including co-actuation of neighboring elements^16,71–73^, could complement FiberTron by resolving local heterogeneity averaged out during pileup, and augment training data. Furthermore, pre-processing Fiber-seq data with read-level classifiers, e.g., distinguishing tumor-*vs*. normal-derived reads^74^, is likely to improve performance on bulk samples. Third, conditioning on observed methylation or accessibility helps where epigenetic state rather than sequence drives allelic imbalance, without the need for additional assays^20^. Likewise, integrating direct RNA-seq information on transcript splicing (e.g., junction read counts), polyadenylation, or base modifications would enable models to capture additional biology on post-transcriptional regulation. Fourth, replacing self-attention by other blocks may not only increase computational performance but also accuracy. An example is Hydra, a bidirectional state-space block in which complexity scales linearly with sequence length, rather than quadratically^21,75^. Finally, single-sample models complement population-scale approaches that pair long-read genomes with single-cell atlases across many donors, combining base resolution and single-molecule linkage with statistical power^76^.

In summary, we demonstrate that small S2F models, trained on multi-omic long reads from a single sample, can match large compendium-trained models such as Borzoi and AlphaGenome. Trained FiberTron instances predict long-range methylation, accessibility and transcript coverage at base resolution from reference sequence alone, and allow variant-effect predictions to be tested against the samples’ own haplotypes. This brings sequence-based interpretation of non-coding and structural variation within reach of individual labs, while addressing clinically relevant biases and gaps in consortium data, such as rare tumors and individual patients.

## Methods

### Ethics approval and patient consent

The Royal National Orthopaedic Hospital (RNOH) Biobank is approved by the National Research Ethics Committee of the Health Research Committee (reference: Integrated Research Application System (IRAS) project identifier: 272816). This study was approved by the UCL/UCLH Biobank for Studying Health and Disease – EC39.25 The Multi-Omic Diversity of Clear Cell Sarcoma. Written informed consent was obtained from the individual participant.

### Data acquisition

To showcase FiberTron’s capabilities, we analyzed data from a human lymphoblastoid cell line (HG002, GM24385/NA24385) from the personal genomes project^77^, a human melanoma cell line (MM001), and a rare clear cell sarcoma patient sample (CCS15).

#### Sample information

The HG002 (GM24385) lymphoblastoid line was obtained from the Coriell Institute for Medical Research. The MM001 melanoma line was derived from a patient biopsy by the Laboratory of Oncology and Experimental Surgery (Prof. Dr. G. Ghanem) at the Institut Jules Bordet, Brussels and has been described previously^6^. For CCS15, fresh-frozen tissue from a metastasic lesion was obtained from the Stanmore Musculoskeletal Biobank, a satellite of the UCL/UCLH Biobank (HTA Licence Number 12055), which was approved by the National Research Ethics Committee (reference 20/YH/0088).

#### Cell culture & tissue processing

HG002 cells were cultured in RPMI1640 medium (Gibco, 21875034) and supplemented with 15% fetal bovine serum (FBS; Biowest, S1810-500). MM001 was cultured in Ham’s F-10 Nutrient Mix (Invitrogen) + 10% FBS (Invitrogen). For both HG002 and MM001, cell harvesting was performed by gentle pipetting, pelleting at 300 *×* g for 5 min at 4 ^*°*^C, and washing twice with PBS. Fresh-frozen CCS15 biopsy tissue was sectioned into 15 curls of 20 *µ*m thickness using a cryostat. All curls were collected in a 1.5 mL Eppendorf tube and re-suspended in 500 *µ*L of ice-cold nuclei extraction buffer (10mM Tris-HCl pH 7.4, 10 mM NaCl, 0.1% BSA, 0.5 mM spermidine pH 7.4, 0.1% Tween-20, 0.02% digitonin). The mixture was transferred to a pre-chilled KIMBLE dounce tissue grinder and homogenized on ice with 10–20 strokes of pestle “A” until complete tissue disruption was achieved.

#### Fiber-seq nuclei extraction & open chromatin labeling

Nuclei extraction was performed on 4 million viable cells (for both HG002 and MM001) by re-suspending the cell pellet in 500 *µ*L of the same ice-cold nuclei extraction buffer. The tissue homogenate was incubated for 4 min on ice, while the cell homogenates were incubated for 3 min on ice. Both were quenched by addition of 1 mL ice-cold Wash Buffer (10 mM Tris-HCl pH 7.4, 10 mM NaCl, 0.1% BSA, 0.5 mM spermidine pH 7.4, 0.1% Tween-20). The tissue homogenate was filtered through a 100 *µ*m cell strainer, and the cell homogenates through a 40µm cell strainer. The nuclei were pelleted at 300*×*g for 5 min at 4°C. The nuclear pellet was washed twice by re-suspension in 500 *µ*L Wash Buffer and re-centrifugation, followed by a final filtration through a 40 *µ*m cell strainer. Nuclei were collected in a 2 mL Eppendorf tube, counted, and quality-checked using a LUNA-FL^−^ Automated Fluorescence Cell Counter (Logos Biosystems).

Nuclei were subjected to 6mA labeling as follows. A total of 1*×*10^6^ nuclei per sample were pelleted by centrifugation at 300 *×* g for 5 min at 4 ^*°*^C. The pellet was gently re-suspended using wide-bore tips in 200 *µ*L Fiber-seq labeling buffer (15 mM Tris, pH 8.0; 15 mM NaCl; 60 mM KCl; 1 mM EDTA, pH 8.0; 0.5 mM EGTA, pH 8.0; 0.1% BSA; 0.5 mM spermidine, pH 7.4), freshly supplemented with 0.8 mM S-adenosylmethionine (NEB, B9003S) and 3.5 *µ*M recombinant Hia5 N6-adenine methyltransferase, purified as described previously^14^. The mixture was incubated in a thermomixer for 10 min at 25 ^*°*^C and 800 rpm. The reaction was stopped by proceeding directly to DNA extraction using the Monarch High Molecular Weight (HMW) DNA Extraction Kit for Tissue (NEB, T3060).

#### PacBio Fiber-seq library preparation

Fiber-seq libraries were prepared for the PacBio Sequel IIe (HG002 and MM001) and PacBio Revio platforms (CCS15). For all samples, extracted, 6mA-labeled HMW DNA was utilized as the starting material. Specifically, 3 *µ*g for HG002 and MM001 and 2 *µ*g for CCS15 of this 6mA-labeled HMW DNA was diluted in 130 *µ*L low TE buffer and sheared to approximately 20 kb. Shearing was performed using a Megaruptor 3 (Diagenode, B06010003) equipped with Megaruptor 3 Shearing syringes (Diagenode, E07010003) at a shearing speed of 31. The resulting sheared DNA was purified using a 1X SMRTbell cleanup (PacBio) and eluted in 47 *µ*L of elution buffer. For all samples, SMRTbell library construction (end repair and A-tailing, adapter ligation and nuclease treatment) was performed using the SMRTbell Prep Kit 3.0 (PacBio, 102-182-700) according to the manufacturer’s instructions. Specifically, for CCS15 an extra size selection was performed using a Pippin HT system (SageScience) with HT 0.75% gel (Sopachem, HPE7510), selecting 6mA labeled HMW DNA fragments starting from 12 kb with a 30 min elution time to yield gDNA fragments of 12 to 40 kb. The size-selected library underwent a 1*×* SMRTbell cleanup and was eluted in 25 *µ*L elution buffer.

For HG002 and MM001 polymerase binding was performed using the Sequel IIe binding kit 3.2 (PacBio, 102-333-300) according to the manufacturer’s instructions. Libraries were sequenced on a PacBio Sequel IIe using three SMRT Cells 8M (PacBio, 101-389-001), to a combined yield of around 100 Gb of HiFi reads per library. For CCS15, the Revio SPRQ polymerase kit (PacBio, 103-520-100) was used according to the manufacturer’s instructions. The final library was sequenced on a PacBio Revio instrument using one Revio SMRT Cell (PacBio, 102-202-200).

#### RNA extraction & ONT direct RNA-seq library preparation

Cell line material was pelleted prior to RNA isolation. Tissue from CCS15 was pulverized into a fine powder using a mortar and pestle on dry ice. Total RNA from HG002, MM001 and CCS15 was isolated using the Maxwell RSC simplyRNA Cells Kit (Promega, AS1390) according to the manufacturer’s protocol. RNA concentrations were measured using the Qubit RNA HS Assay Kit (ThermoFisher, Q32852). Direct RNA sequencing libraries were prepared using the Oxford Nanopore Technologies (ONT) Direct RNA Sequencing Kit (SQK-RNA004) according to the manufacturer’s protocol.

Briefly, 1000 ng, 1800 ng and 1000 ng of total RNA of HG002, MM001 and CCS15 respectively, were adjusted to 8.5 *µ*L with nuclease-free water. The RNA Calibration Strand component was excluded from the libraries. The RNA was ligated to the RT Adaptor (RTA) using T4 DNA Ligase (NEB, M0202M) in NEBNext Quick Ligation Reaction Buffer (NEB, B6058) for 10 min at room temperature. Reverse transcription was performed using Induro Reverse Transcriptase (NEB, M0681) by incubating at 60 *°*C for 30 minutes and 70 *°*C for 10 minutes. The resulting RNA/cDNA complex was purified using RNAClean XP beads (Beckman Coulter, A63987). The RNA Ligation Adapter (RLA) was then ligated to the RNA/cDNA duplex using T4 DNA Ligase (NEB, M0202M) for 10 min at room temperature. Following a second bead cleanup with RNAClean XP beads and two washes with Wash Buffer (WSB), the final library was eluted in 33 *µ*L RNA Elution Buffer (REB) at room temperature for 10 min. The eluted libraries were sequenced according to ONT protocols on a PromethION 2 Solo platform. Each library was sequenced on one PromethION Flow Cell (ONT, FLO-PRO004RA) using standard RNA sequencing run parameters.

#### Reference genome and genomic annotations

Input DNA sequences were extracted from the Telomere-to-Telomere-complete CHM13v2.0 human reference genome^12^. Functional modality targets were extracted from the in-house generated PacBio HiFi Fiber-seq and ONT direct RNA-seq datasets. Gene annotations used for model evaluations were based on JHU RefSeq + Liftoff v5.1 annotations (https://github.com/marbl/CHM13), unless otherwise specified.

### Data pre-processing

Data pre-processing was performed using custom-developed Nextflow (v.24.10.3)^78^ pipelines. For raw unaligned Fiber-seq BAM files (*≥*30*×* HiFi coverage) the pipeline performs modified basecalling (6mA modification only) and Fiber-seq Inferred Regulatory Element (FIRE) calling with fibertools (v.0.6.2)^79^, alignment with minimap2 (v.2.28)^80^ and sorting, indexing and merging of the resulting BAM files with samtools (v.1.21)^81^.

For raw direct RNA-seq POD5 data the pipeline performs basecalling using dorado (v.0.9.1)^82^. Then a modified pipeline based on nf-core/nanoseq (v.3.1.0)^83^ performs alignment with minimap2 (v.2.17) and transcript quantification with stringtie2 (v.2.1.4)^84^ and featureCounts (v.2.0.1)^85^. The pipeline emits an expression count matrix and stranded BIGWIG files generated with ucsc_bedgraphtobigwig (v.377)^86^.

### Training/validation/test data generation

All chromosomes, excluding chrY and MT, were randomly split into training, validation and test sets to minimize data leakage between partitions (test: chr2, chr9; validation: chr13, chr14; training: all others). Chromosomes were fragmented into overlapping regions of 524288 bases (524 kb) in length with an overlap of 65536 bases (65 kb). The sampled regions were filtered based on coverage depth, with regions being included if and only if they have a coverage in the range [4, median coverage + 5*×*MAD], where MAD is the genome-wide median absolute deviation in the coverage. Genome wide coverage statistics were computed using mosdepth (v.0.3.3)^87^.

To further increase the implicit size of the training data without introducing additional memory overhead, online data augmentation (during batch loading) was applied. Only data augmentations that preserve the biological meaning of the input DNA sequences and that of the corresponding modalities were applied. Specifically, training batches were randomly augmented by applying the following transformations retaining the relationship between input and targets:

- Left or right shift of input sequence and target boundaries. Shift size was randomly sampled from the range [1, 16384].
- Reverse complement of input sequences and targets. For stranded targets, the forward and reverse strands were also exchanged.

#### DNA sequence formatting

DNA sequences of regions passing the coverage depth filter were extracted from the CHM13v2.0 human reference genome FASTA file and one-hot encoded.

#### Fiber-seq data formatting

Base resolution CpG methylation and chromatin accessibility information was obtained by extracting the ML tag values from the Fiber-seq BAM file using samtools (v.1.21) mpileup. The ML tag is an optional field tag of SAM/BAM files that contains the probability of a base being modified in a read. ML tag values belong in the discrete range *A* = [0, 255] with each integer value *n* corresponding to [*n/*256, (*n* + 1)*/*256) probability range.

First, the ML tags of 5mC and 6mA modifications, for read-level CpG methylation and chromatin accessibility respectively, were binarized:

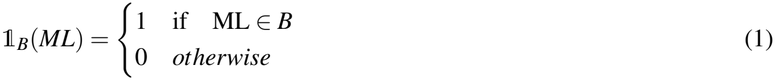

where *B* ⊂ *A, B* = [128, 255]. Then for every base of the 524 kb reference vectors a modification probability percentage in the range [0, 100] was computed reflecting the proportion of 5mC or 6mA modified reads at that base.

Both vectors were then masked to fully comply with the reference sequence, meaning that 5mCpG calls at non-reference CpG sites or 6mA calls at reference G or C bases (i.e., typically at sample-specific polymorphisms) were set to 0 modification probability.

Chromatin accessibility signals were next smoothed and interpolated. Specifically, percentage modification probabilities were linearly interpolated to guanine and cytosine bases from the nearest adenine or thymine base and then smoothed using a rolling moving average with a 5 bp window.

Fiber-seq Inferred Regulatory Elements (FIRE) counts per base were computed using fibertools (v.0.6.2) pileup command with 128 set as the minimum ML tag threshold. Then every base of the 524 kb reference vectors was given a percentage modification probability in the range [0, 100] depending on the FIRE count/total coverage fraction of each base.

#### Direct RNA-seq data formatting

For each reference region forward and reverse strand gene expression vectors were generated by extracting the base resolution raw coverage from the generated BIGWIG files. As in Linder *et al*.^9^, to account for the relatively large dynamic range of RNA-seq and to limit the contribution of highly expressed genes during model training, the raw coverage was normalized by exponentiation of the base-resolution coverage by 3/4. If the exponentiated values were larger than 384, an additional square-root transformation was applied on the residual value:

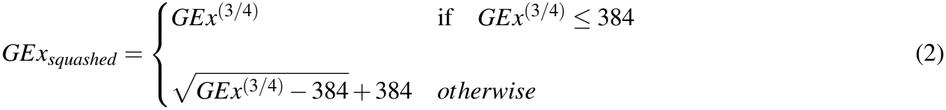

### FiberTron model architecture

The FiberTron architecture is inspired by SoTA deep neural network architectures from the field of genomics^4,5,8,9,88^. The model has a hybrid convolutional transformer U-net architecture^9^, consisting of an encoder part which downsamples the input data and a decoder part that upsamples the data back to base-resolution. The model was developed using the PyTorch^89^ deep learning ecosystem: torch (v.2.5.1), torchmetrics (v.1.6.1), einops (v.0.8.0).

#### Encoder

The encoder consists of an encoding block and a series of downsampling blocks that extract local sequence patterns using convolutional layers and perform two-fold reduction of the input spatial dimension (sequence length) by utilizing pooling operations^90^. Pooling layers are followed by a root mean square normalization layer^91^ and GELU^92^ activation functions. These downsampling blocks are applied until the sequence is aggregated to features corresponding to 512 bp. From the 16 bp aggregation level and onward, the downsampling blocks are followed by transformer encoder blocks that allow to capture contextual information across sequence positions. Each transformer encoder block consists of three components: the input projection component (projects input embedding into query, key and value tensors), self-attention mechanism^93^ with rotary position embeddings (RoPE)^94^ and a feed-forward network with a gated linear unit^95^. A key difference with existing applications is that the self-attention mechanism is interleaved between the convolutional downsampling blocks. Instead of operating at a single resolution of the input data, the transformer blocks can hence operate at multiple levels of the input sequence data, enabling FiberTron to extract coarser but also more fine-grained features to model interactions between distal regulatory elements.

#### Decoder

The decoder consists of upsampling blocks, with every block doubling the output embedding spatial dimension and decreasing the aggregation two-fold until the embedding reaches base-pair resolution. At each different resolution level the upsampling output embeddings are enriched with the intermediate feature maps from the corresponding downsampling blocks and then passed through a separable convolution layer.

#### Multi-task head

Regions located near the start and end of an input sequence are at a relative disadvantage compared to the sequence center as the model can only observe part of their regulatory context^8^. To address this, the base-resolution output embedding from the encoder part is passed to a cropping layer that trims 98,304 bases from either side. Trimming the ends of the embedding ensures that each sequence position for which the loss is computed, has at least *≈* 98kb sequence upstream and downstream. The trimmed embedding then passes through a block that performs a point-wise operation and finally through the multi-task output head that performs a convolution operation followed by a softplus activation function to ensure the positivity of the predictions.

### Training framework

The model is trained in a multi-task supervised setting to predict the processed Fiber-seq and direct RNA-seq-derived functional modalities from the one-hot encoded reference DNA sequences as input. More specifically, the model predicts base-resolution DNA methylation at CpG sites, genome wide chromatin accessibility and stranded full-length gene expression coverage.

The aforementioned modalities follow distributions with different statistical properties (akin to Beta, Gaussian, and Poisson distributions respectively). To account for the latter and avoid loss scaling differences, a weighting scheme based on task-specific homoscedastic uncertainty^96^ was utilized. This multi-task loss function allows to combine multiple different loss functions and alleviates the manual finetuning of task loss weights as weighting is learned during model training procedure. For CpG methylation and chromatin accessibility modalities the mean squared error (MSE) loss function was used whereas for gene expression a custom Poisson Multinomial Negative Log Likelihood (PMNLL)^9^ loss was used.

Model instances were trained on 4 NVIDIA H100 GPUs on the Flemish Supercomputer Center (VSC) HPC cluster using PyTorch’s Distributed Data Parallel module. Batch size was set to a total of 24 sequences (6 per GPU; maximum allowed number not exceeding the 80 GB RAM of each GPU). Adam optimizer with decoupled weight decay regularization^97^ was used with an initial learning rate of 1 *×* 10^*−*3^. A linear learning rate warm-up schedule^98^ was applied for the first 5,000 training iterations. For the remainder of the training procedure, the learning rate was dynamically adjusted using cosine annealing with warm restarts^99^. Gradients were clipped to a maximum global norm of 0.2^8^. Early stopping was applied if for 20 consecutive training epochs no metric improvement was observed on the validation set. The metric tracked during model training was the Concordance Correlation Coefficient (CCC)^100^. Unlike Pearson’s *r*, CCC penalizes deviations in location and scale from the identity line, so it reflects both precision and accuracy (bias) of the predictions.

### VCF file liftover

Liftover of VCF coordinates between reference genome assemblies was performed using GATK (v.4.1.6)^101^ LiftoverVcf (Picard; v.3.3.0)^102^ with default parameter values. This lifts over variants that map exactly from GRCh37/GRCh38 to T2T-CHM13v2.0 but does not recover variants with swapped reference and alternative alleles. To recover variants with swapped reference/alternative alleles, LiftoverVcf was run again, with the RECOVER_SWAPPED_REF_ALT flag. Notably, this feature does not recover multiallelic variants, so to recover these variants, we first separated them into multiple biallelic variants, performed liftover using the RECOVER_SWAPPED_REF_ALT tag, and converted these back into their multiallelic representations.

### Variant calling and phasing

For HG002, SNVs and indels were generated using Clair3 (v.1.0.9)^103^ and SVs were called using Sniffles (v.2.5.2)^104^, respectively. For MM001 and CCS15 SNV and indel calls were generated using ClairS-TO (v.0.4.0)^105^ and SV calls with Severus (v.1.2)^106^. Resulting variants were phased using LongPhase (v.1.7.3)^107^.

### Variant effect scoring metrics

The effect of genetic variation on the different functional modalities was quantified using the following variant effect scoring approaches. Per notation, *L* is the length of the focus window; *m* ∈ {1, 2, 3, 4} denotes the CpG methylation, chromatin accessibility, and gene expression on the plus and minus strand modalities respectively; *i* the base index and *y*^*re f*^ = *F*(*x*^*re f*^) and *y*^*alt*^ = *F*(*x*^*alt*^) ∈ ℝ^*L×m*^. *ρ*_*m*_ and 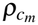 are the Pearson’s and Concordance correlation coefficients respectively.

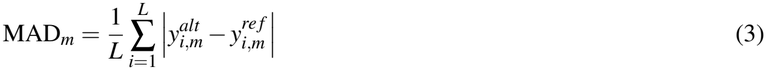

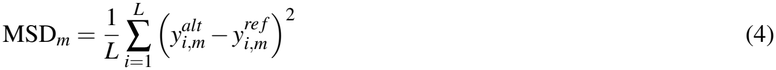

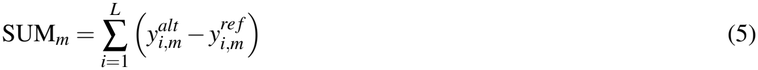

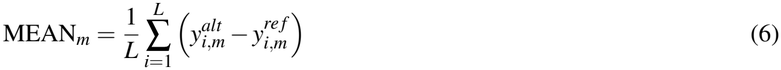

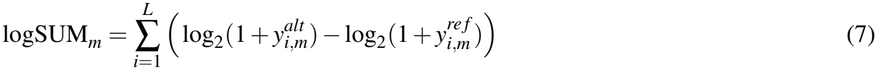

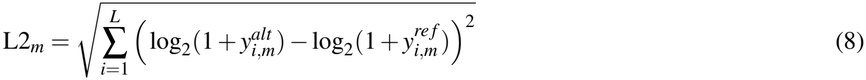

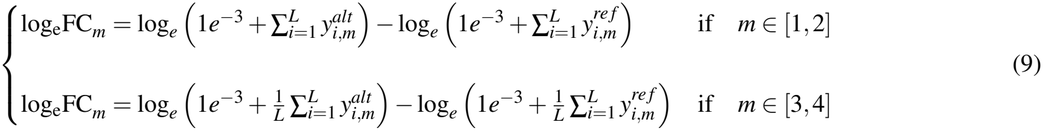

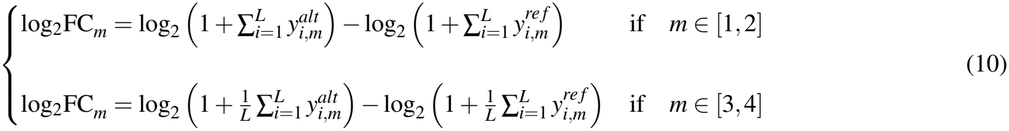

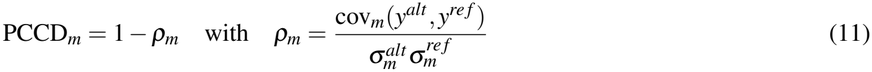

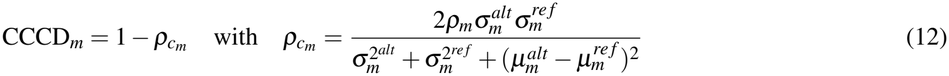

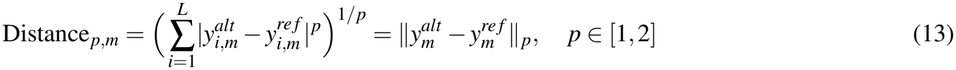

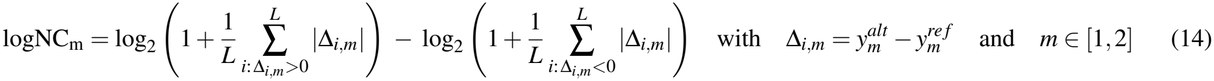

For CpG methylation and chromatin accessibility, variant effects were assessed in a window centered around the variant ranging from 125 bp to 2 kb. For gene expression, variant effects were aggregated over all annotated gene exons falling inside FiberTron’s output window. For eQTL data, gene expression variant effects were computed for the annotated gene paired with each variant.

### Complex variant effect scoring strategy

Compared to SNV, scoring indels and SVs poses several technical challenges. Evidence suggests that sequence shifts introduce technical variance in convolutional S2F architectures through the misalignment of pooling blocks and output boundaries^108^. To address this, we adopted a scoring strategy in which, for every variant, four different sequences for the ALT allele were generated using data augmentation shifts. Two which preserved alignment on the left- and right-most sides of the REF genomic region^108^, respectively, and two that preserved alignment of the center with excess padding either distributed on the left or right side (Supplementary Figure S1a-b). For consistent positional comparison, the resulting predictions for each sequence (REF and four augmented ALT sequences) were aligned back to a common coordinate space (expanded with zero-padding) and a binary exclusion mask was generated to account for missing bases in REF or ALT sequences in the case of insertions and deletions, respectively (Supplementary Figure S1c-d). On the basis of our empirical results, the mean of the left- and right-anchored constructs yielded the highest performance metrics for causality prediction tasks, whereas the center-anchored constructs exhibited greater concordance with the ground-truth effect size comparisons.

### eQTL prediction tasks

For eQTL prediction evaluations, the Sum of Single Effects regression (SuSiE)^109^ fine-mapped Genotype-Tissue Expression (GTEx) Portal (v.8) re-processed dataset^44^ was utilized. Only variants with a posterior inclusion probability (PIP) *≥* 0.9 in lymphocytes (“EBV-transformed lymphocytes” annotation) were retained, resulting in a set of 153 putative causal variant *×* protein-coding gene pairs. Matched putative non-causal variants were sampled from the HG002 set of heterozygous SNVs/INDELs accounting for distance to the nearest TSS and expression level of the corresponding gene, yielding 152 putatively non-causal variants. All variants passed the sequencing depth filtering threshold applied during data modality generation. To make predictions robust to incidental properties like exact bin alignment and input orientation, test-time augmentation (TTA) by reverse complementing^10^ was applied.

### eQTL effect size (coefficient) prediction

FiberTron’s ability to predict the effect size of known causal variants was assessed on the GTEx putatively causal eQTLs. While observed (SuSiE *β* coefficients) and predicted (FiberTron variant effect scores) effect sizes may differ in scale, agreement is expected in their relative ranking and direction (sign). To quantify concordance between observed and predicted effect sizes, we computed and reported Spearman’s and Pearson’s correlations.

### eQTL effect direction (sign) prediction

eQTLs were classified according to their effect on gene expression, with cases having *β ≥* 0 designated as “positive” (n=71) and cases with *β <* 0 as “negative” (n=82). To evaluate whether FiberTron can predict whether a causal variant has a positive or negative effect, we computed the auROC for the binarized FiberTron-derived scores against the binary GTEx eQTL sign labels.

### eQTL causality prediction

The zero-shot ability of FiberTron to differentiate causal from non-causal eQTLs was evaluated by computing auROC for the FiberTron-derived scores against the binary GTEx eQTL causality labels.

### Splicing variant effect predictions

Using genomic loci with known *cis*-directed splicing events in lymphoblastoid line GM12878^45^, we investigated whether FiberTron can pinpoint causal variants and predict their transcriptional effects. Since FiberTron models are not explicitly trained with splice junction information data, causal variants are identified using variant effect scores on the gene expression modality. Two scores are computed: a gene-specific score corresponding to the affected gene and an exon-specific score corresponding to the exons closest to the candidate causal variant.

### *De novo* allele-specific event identification

Allele-specific differential methylation (DMR) and chromatin accessibility (DAR) calling was done using modkit (v.0.5.0)^32^. Loci with allele-specific differential methylation or chromatin accessibility patterns were identified using two approaches: (a) by providing modkit with a set of genomic regions of length *L* centered around called heterozygous variants, or (b) using modkit’s (fine-grained) segmentation algorithm and intersecting the identified segments with the heterozygous variants. The resulting differential methylation and accessibility calls can be stratified into small, medium or large effect size calls using the following thresholding on the modkit-computed Cohen’s *h* statistic:

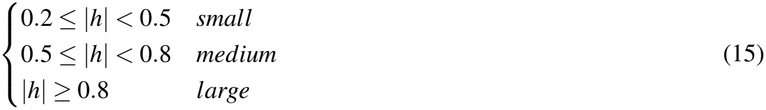

Allele-specifically expressed (ASE) genes were identified using longcallR (v.2.0.0)^110^. High-confidence calls were obtained by requiring identified genes to be supported by both longcallR transcriptome-inferred heterozygous variants and ground truth WGS-based phased variant calls. The resulting high-confidence ASE genes were further filtered by retaining only those with a Benjamini-Hochberg (BH)-corrected binomial test p-value of *≤* 0.05. (*H*_0_ : *π* = 0.5; *H*_1_ : *π* ≠ 0.5).

### ASCAV prediction tasks

Utilizing previously identified allele-specific chromatin accessibility variants (ASCAVs) in melanoma line MM001^6^, we investigated whether FiberTron can differentiate causal from non-causal chromatin accessibility variants.

### Zero-shot ASCAV causality prediction

FiberTron-derived scores for ASCAVs (n=1,840) were compared to a background score distribution generated from heterozygous MM001 SNVs (called using ClairS-TO v.0.4.0)^105^, overlapping known cREs, and not having an SV in the same 524 kb window (n=20,807). FiberTron’s ability to distinguish causal from non-causal ASCAVs was assessed at different false-positive rates (FPR) for all variant effect scoring metrics as described in Methods. Fisher’s exact test was used to test association in the 2 *×* 2 contingency table at FPR = 0.05.

### Fusion gene expression prediction

Using the phased heterozygous SV calls generated for CCS15, we reconstructed the exact sample-specific *EWSR1::ATF1* fusion gene sequence and its reciprocal *ATF1::EWSR1*. These reconstructed sequences were subsequently used as custom references for re-alignment of CCS15 direct RNA-seq reads to establish the ground-truth fusion profiles, and as input to FiberTron for prediction of the corresponding expression profiles.

### Supervised variant pathogenicity prediction

The Clinical Significance of Human Genetic Variants (ClinVar) database (release of 30 May 2026; v.4.1) was downloaded from https://ftp.ncbi.nlm.nih.gov/pub/clinvar/vcf_GRCh38/weekly/ and lifted over from GRCh38 to the T2T-CHM13v2.0 assembly. Variants were retained based on the following criteria: (a) curation review status of more than one gold star, (b) “Benign” or “Pathogenic” | “Likely_pathogenic” | “Pathogenic/Likely_pathogenic” clinical significance annotation, (c) molecular consequence annotation falling in the following set: {“intron_variant”, “synonymous_variant”, “missense_variant”, “nonsense”, “splice_donor_variant”, “splice_acceptor_variant”, “5_prime_UTR_variant”, “3_prime_UTR_variant”}, (d) “likely_benign” AlphaMissense^111^ annotation, and (e) autosomal. Retained variants were divided into six categories based on their molecular consequence annotation and distance to the nearest splice site^10^, designated as “Intronic”, “Synonymous”, “Splicing”, “Missense”, “Nonsense”, and “UTR”.

To investigate whether FiberTron can distinguish pathogenic from benign variants, a Random Forest (RF) classifier (scikit-learn v.1.6.1^112^ algorithm implementation) was trained using FiberTron variant effect scores as input. Variants were split into training and test sets stratified by pathogenicity label and annotation category. The binary classifier was trained and evaluated using a 10-fold cross-validation framework. To evaluate the performance of the classifier, we computed average auROC and auPRC scores across the held-out test folds.

### Supervised variant rarity prediction

As rare variants are enriched in pathogenic alleles compared to common variants, we next investigated whether FiberTron could distinguish these rare from common variants. To this end, a set of 22,201 rare and 22,246 matched common (allele frequency > 0.05) biallelic SNVs was obtained from the Genome Aggregation Database (gnomAD)^49^ release v.3.1.2 lifted over to T2T-CHM13v2.0 (original source https://gnomad.broadinstitute.org/downloads) as downloaded from https://github.com/marbl/CHM13.

Variant sampling was restricted to genomic regions overlapping known cREs, while excluding variants located within CpG islands, protein-coding gene exons, and low-complexity regions (including repetitive elements, centromeres, and telomeres). In addition, sampled common variants were further constrained to those sharing the same tri-nucleotide sequence context as their matched rare variant. For all variants in the obtained sets, CADD (v.1.7)^50^, PHRED, and RAW scores were compared to FiberTron-derived scores.

Three RF classifiers (scikit-learn v.1.6.1^112^) were implemented using as input features; (a) FiberTron-derived scores capturing direction, total signal and profile, (b) CADD (v.1.7), PHRED, and RAW scores and (c) all scores from (a) and (b). The binary classifiers were trained and evaluated using a 10-fold cross-validation framework (variants were split into training and test sets stratified by rarity label). The average Precision, Recall, F1-score, and auROC across held-out test folds were computed and used to evaluate the classifiers.

### Attention matrix processing & visualization

Higher-order chromatin structures and long-range interactions learned by FiberTron may be extracted from the attention weight matrices of the hierarchical self-attention layers. As the different attention heads tend to capture the same trends, the mean attention of all four heads was computed for each self-attention layer. We observed that, at each resolution, the attention mechanism captured distinct patterns. Therefore, to utilize all captured information at the finest resolution (16 bp), the attention weight matrices of the coarser resolutions were up-scaled to 16 bp (nearest neighbor interpolation), and re-normalized so each row summed to 1.0. For visualization of the complete attention maps, intensity values were clipped at the 0.5% and 99.5% percentiles of the empirical distribution. For visualization of zoomed-in regions of the attention maps, values were clipped at the 0.01% and 99.9% percentiles. To attenuate high-frequency noise in the zoomed-in attention maps, a Gaussian filter with standard deviation *σ* =0.5 and truncation parameter set to 2.0 was applied. All attention weight matrices were initially averaged over the reverse-complemented input sequences.

### Enhancer × gene pair interaction detection

Utilizing a set of validated enhancer-gene interaction pairs in lymphoblastoid line GM12878^43^, we systematically assessed whether FiberTron assigns greater attention to these loci relative to their corresponding genomic context. For each gene in the set belonging to the held-out test chromosomes, the attention weight matrix was computed as described before (without clipping). For each annotated enhancer–gene pair, we extracted the slice of the attention map corresponding to the attention directed from the gene promoter to the enhancer. To determine whether systematic differences in attention magnitude were detectable, gene promoter sequences were shuffled in silico (n = 10). Differences in attention between the wild-type and shuffled sequences were evaluated using a binomial sign test applied globally across enhancer-gene pairs.

### DNA sequence attribution

Although numerous explainability algorithms have been developed, interpreting S2F DL genomic models and extracting biologically meaningful insights remains non-trivial. To derive mechanistic insights (e.g., known sequence patterns, TF recognition motifs) from FiberTron’s predictions, different *post hoc* attribution methods were applied. Methods utilized were either based on gradient saliency, which is computationally efficient but more noisy (off-simplex gradient noise^113^), or on in silico mutagenesis (ISM), which provides better-calibrated attributions, but becomes intractable for longer sequences.

Let *f* represent a FiberTron model instance and *s* the input sequence. In gradient-based (saliency) attribution methods, position-specific importance scores of each nucleotide are quantified by calculating the partial derivative of the model’s output with respect to the input 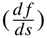. This can be interpreted as performing a first-order Taylor expansion (a linear approximation of *f* at input sequence *s*) where gradients serve as the coefficients of the linear function. The magnitude of these coefficients in absolute value is used to measure feature importance.

ISM attribution scores are derived by systematically perturbing each position in the input sequence and observing the difference in model predictions. The effect of mutation at position *l* from base *b*_0_ to *b*_1_ is derived as follows:

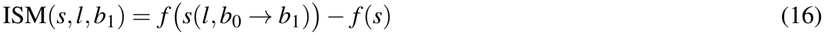

Taylor expansion approximation provides a direct connection between ISM and gradient-based attribution methods^34^. The 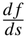 gradient can be interpreted as a finite-difference corresponding to a single nucleotide substitution from *b*_0_ to *b*_1_ at position *l*. Under a first-order Taylor expansion, ISM reflects the linear effect of this substitution on the model output. Conversely, this relationship allows computationally expensive ISM scores to be approximated using gradients:

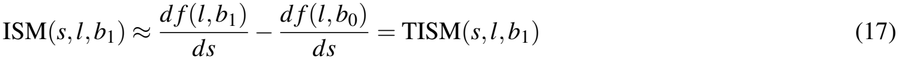

However, as a local linear approximation, TISM may deviate from ISM in strongly nonlinear settings (e.g., activation saturation or redundant regulatory grammar), reflecting a well-established limitation of gradient-based attribution methods^114,115^. Moreover, because the sign of an attribution depends on the selected reference and may be challenging to interpret^116^, we report absolute attribution magnitudes when using TISM, consistent with magnitude-based saliency approaches^117^. This choice identifies positions to which the model is sensitive, but it does not distinguish nucleotides that increase predictions from those that decrease them.

## Data analysis and visualization software

Data analyses and visualization were performed using Python v.3.12.8 (https://www.python.org/), Pandas v.2.2.3 (https://github.com/pandas-dev/pandas), SciPy v.1.15.0 (https://github.com/scipy/scipy), Scikit-learn v.1.6.1 (https://github.com/scikit-learn/scikit-learn), NumPy v.1.26.4 (https://github.com/numpy/numpy), PyTorch v.2.5.1 (https://github.com/pytorch/pytorch), Matplotlib v.3.10.0 (https://github.com/matplotlib/matplotlib), seaborn v.0.13.2 (https://github.com/mwaskom/seaborn), tangermem v.1.0.0 (https://github.com/jmschrei/tangermeme), methylartist v.1.5.3 (https://github.com/adamewing/methylartist), and IGV v.3.0.0 (https://github.com/igvteam/igv.js). Borzoi 6503ea5 (https://github.com/calico/borzoi) and AlphaGenome v.0.9.0 (https://github.com/google-deepmind/alphagenome) analyses were performed using Google Colab (https://research.google.com/colaboratory). Schematic representations of the model architecture and complex variant scoring strategy were generated using drawio^118^ (https://github.com/jgraph/drawio).

## Acknowledgements

This work was supported by VIB and KU Leuven Internal Funds ([C14/22/125] SymBioSys) and project grants from the Research Foundation – Flanders (FWO; [G021725N]) and VIB (Grand Challenges [GC06-C05]). It was further enabled by large-scale research infrastructure funding from FWO ([I014324N] T2T-Biology). The compute resources and services used in this work were provided by the VSC (Flemish Supercomputer Center), funded by FWO and the Flemish Government. The Royal National Orthopaedic Hospital provided biobank material for CCS15. The Edward Showler Foundation has generously supported research related to Clear Cell Sarcoma. M.E. is a PhD fellow of the FWO (Grant number: 1176026N). K.D.K. is recipient of a Collen-Francqui research professor mandate and her research is funded by FWO (Grant number: G017226N). N.P. has received funding from Cancer Research UK (Award number 18387) and holds a Cancer Research UK Career Establishment award (RCCCEA-Nov23/100003). L.L. is a postdoctoral fellow of Stichting Tegen Kanker [2025-031]. We are grateful to Andrew B. Stergachis for providing the Hia5 expression construct and the Leuven Genomics Core for their sequencing expertise.

During the preparation of this work, the authors used several large language model (LLM)-based tools for copy editing. Claude (Anthropic; Opus v5.5) was used for copy editing of the code repository accompanying this manuscript, including improvements to code organization, readability and consistency. M365 Copilot (Microsoft; GPT-5), together with Claude (Anthropic; Opus v5.5) and TeXGPT (Writefull, OpenAI; GPT) accessed through Overleaf’s integrated AI features, was used for copy editing of the manuscript text, including improvements to grammar, wording and readability. These tools were not used to generate scientific content, data, analyses, results or conclusions. All LLM-assisted edits to both code and text were reviewed and validated by the authors, who take full responsibility for the content of this publication.

## Author contributions statement

Conceptualization & Methodology: J.D., M.E.; Funding acquisition: J.D., M.E.;

Supervision: J.D., A.S.;

Software (FiberTron implementation): M.E.;

Formal analysis: M.E., L.L.;

Visualization: M.E.;

Data generation: J.V.V., K.T., L.L.;

Resources: J.D., A.S., S.A., K.D.K., N.P., C.D.;

Writing - Original Draft: J.D., M.E.; Writing - Review & Editing: all authors

## Additional information

### Data availability

Sequencing data for CCS15 and MM001 are deposited in the European Genome-phenome Archive (EGA) under study ID EGAS50000002226. The data generated for HG002 have been deposited in the European Nucleotide Archive (ENA) at EMBL-EBI under accession number PRJEB128467.

### Code availability

Code and trained weights will be released under a VIB academic non-commercial license at the latest upon publication and are currently available on request. The source code repository associated with this manuscript, including all analyses and results will be https://github.com/IntGenomicsLab/fibertron_research while model weights and scripts for training, inference, and variant scoring will be at https://github.com/IntGenomicsLab/FiberTron.

### Competing interests

The authors declare no conflicts of interest.

## Extended Data Figures

**Extended Data Figure 1.**
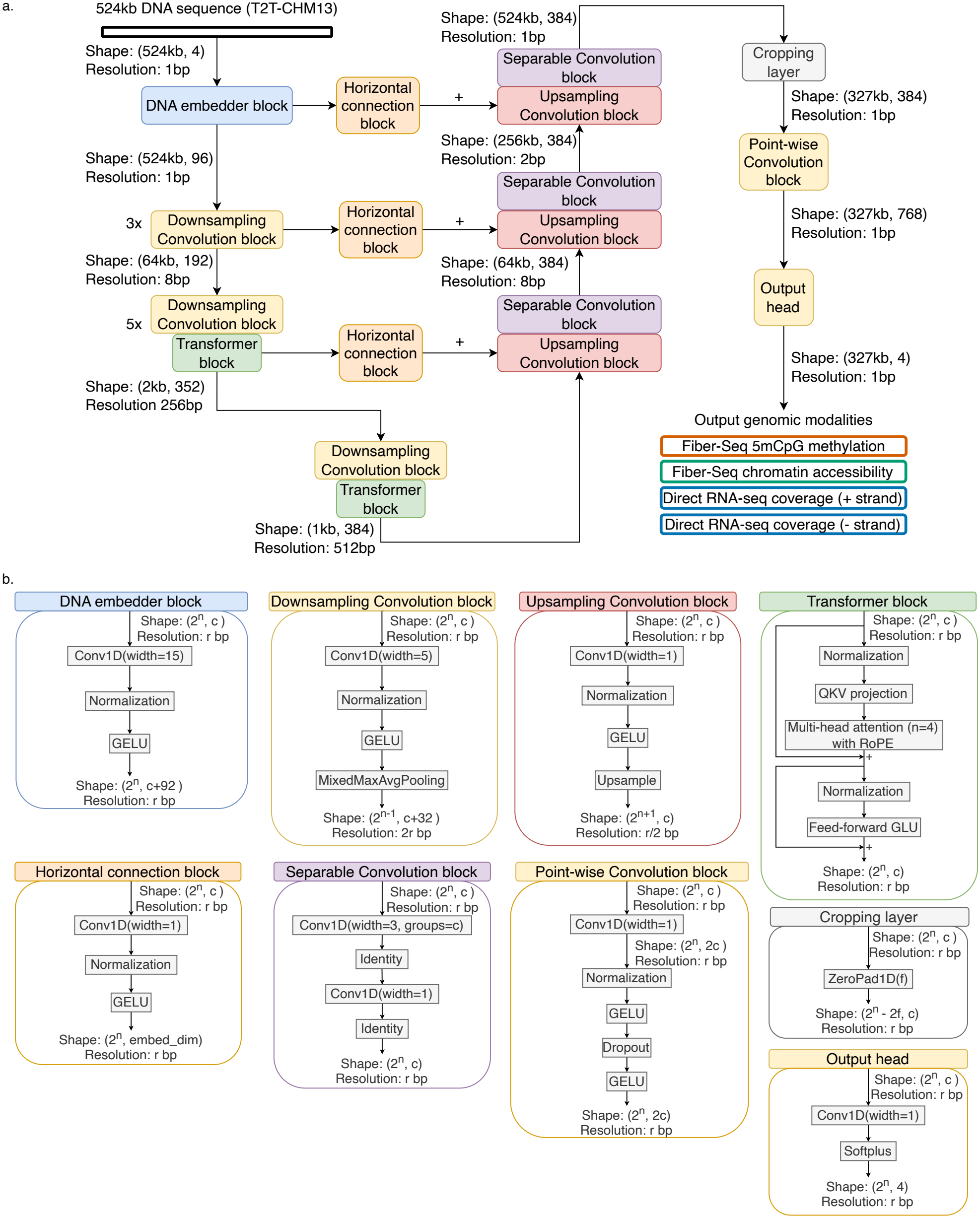
Detailed FiberTron architecture and main component blocks. **a**. Schematic of FiberTron’s architecture highlighting major components and the flow of activations through the network. FiberTron has a U-net backbone^9^ with interleaved self-attention modules at specific resolution levels of the encoder. The decoder path is enriched with intermediate features from the encoder path through residual connections. The resulting feature embedding is symmetrically truncated to guarantee that every predicted genomic position is flanked by a minimum attention context of approximately 100 kb, both upstream and downstream. **b**. Internal structure of the main building blocks employed in model architecture shown in (**a)**.

**Extended Data Figure 2.**
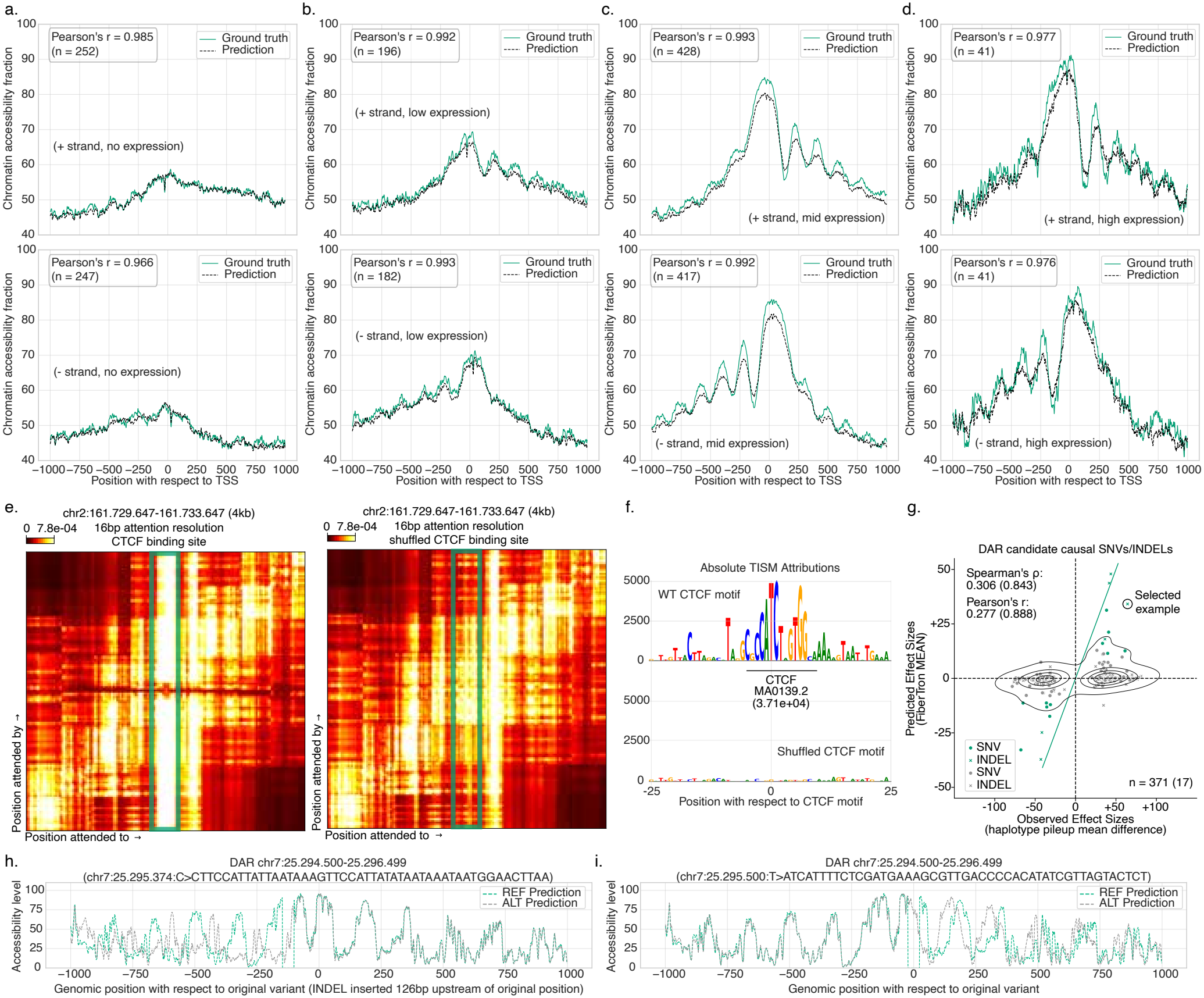
Additional evaluation of chromatin accessibility predictions. **a–d**. Comparative analysis of mean chromatin accessibility profiles at transcription start sites (TSSs) on left out chromosomes, contrasting ground-truth and predicted signals, stratified by gene expression level. (**a**) Silent, (**b**) Low, (**c**) Mid, and (**d**) High. **e**. Magnified view of attention heatmap corresponding to a 4 kb region centered around the example wild-type (left) and shuffled (right) CTCF motif. **f**. Absolute scale TISM^34^ ACWMs and tangermeme^36^ seqlet calling centered around the example wild-type (top) and shuffled (bottom) CTCF motif. **g**. Concordance between Fiber-seq observed allelic effects and FiberTron-predicted variant effect scores at the modkit-identified^32^ allele-specific differentially accessible regions (DARs). Concordance values for candidate causal variants (colored) are provided in parentheses. The highlighted example is circled. **h-i**. FiberTron predictions for two in silico perturbations of the example DAR: (**h**) indel moved upstream of the CTCF binding site and (**i**) replaced with a random, dinucleotide content-matched, insertion of the same length. For **a-f**, all visualizations and reported performance metrics are based on held-out HG002 data.

**Extended Data Figure 3.**
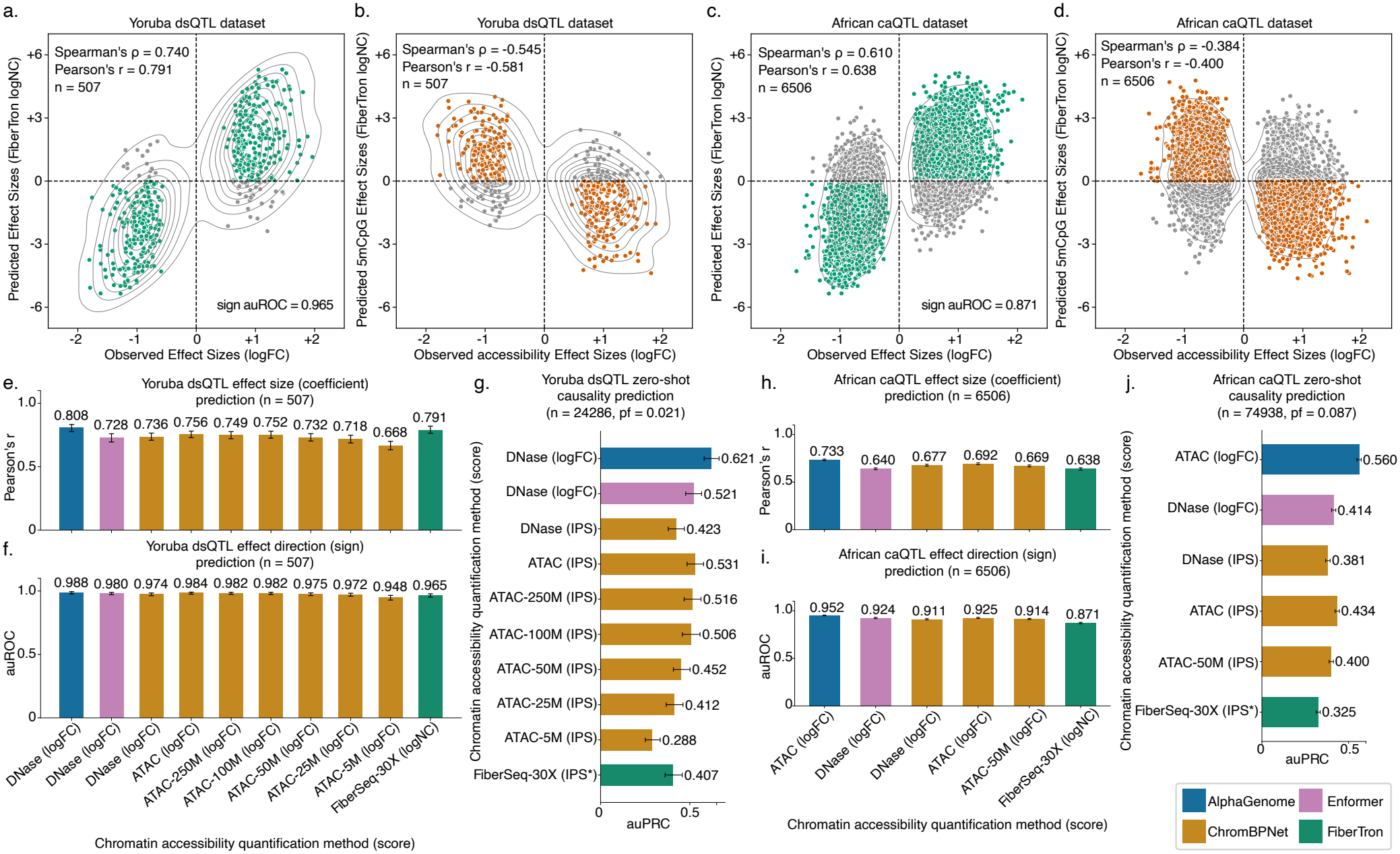
Chromatin accessibility quantitative trait loci (ds/caQTL) prediction tasks. **a**. Comparison between observed and FiberTron-derived effect size (coefficient) of Yoruba ancestry dsQTL dataset. **b**. FiberTron predictions for CpG methylation in relation to the observed chromatin accessibility effect size of Yoruba ancestry dsQTL dataset. **c-d**. Same as (**a-b**) for African ancestry caQTL data. **e-g**. Prediction performance comparison between FiberTron, ChromBPNet^5^, Enformer^8^ and AlphaGenome^10^ models on effect size (coefficient), direction (sign), and causality (zero-shot) of Yoruba ancestry dsQTL data. **h-j**. Same as (**e-g**) for African ancestry caQTL data. Error bars represent bootstrapped 95% confidence intervals (CIs). Abbreviations: IPS^5^, integrative prioritization score; IPS* = MAD *×* CCCD; MAD, Mean absolute difference; CCCD, concordance correlation coefficient difference; logFC, log fold change; logNC, log net change. All visualizations and reported FiberTron performance metrics are based on HG002 data. For ChromBPNet, Enformer and AlphaGenome, scores/predictions were derived from the GM12878 DNaseI-/ATAC-seq tracks.

**Extended Data Figure 4.**
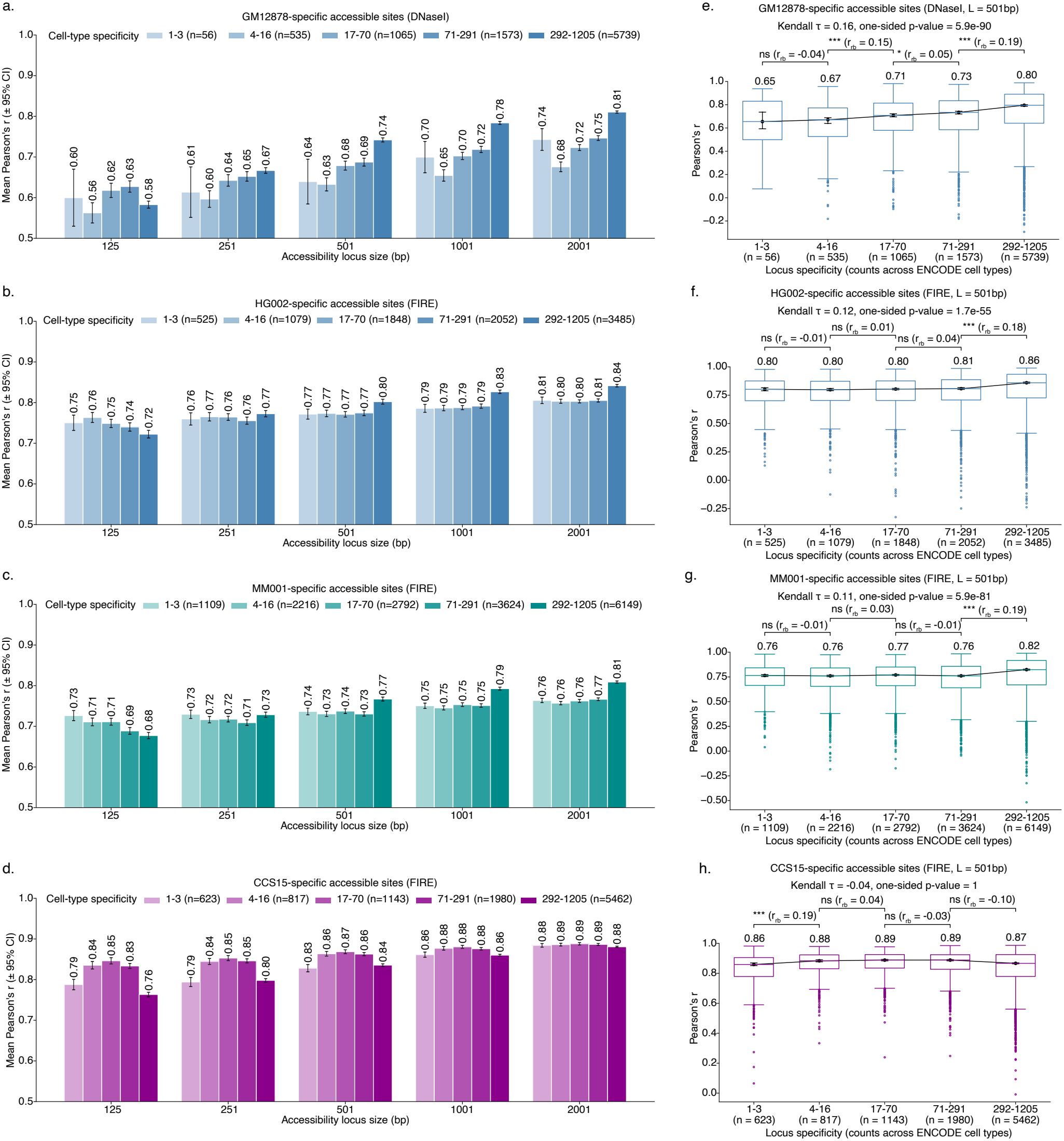
Predictive performance of FiberTron instances for chromatin accessibility at cell-type–specific accessible loci. **a–d**. Mean Pearson’s *r* between ground-truth and predicted Fiber-seq pileup accessibility profiles, evaluated across multiple genomic window spans at cell-type–specific loci: (**a**) GM12878 DNaseI peaks, (**b**) HG002 FIRE peaks, (**c**) MM001 FIRE peaks, and (**d**) CCS15 FIRE peaks. Error bars represent bootstrapped 95% confidence intervals (CIs). **e–h**. Box plots showing Pearson’s *r* computed over a 501 bp window, stratified by cell-type specificity: (**e**) GM12878 DNaseI peaks, (**f**) HG002 FIRE peaks, (**g**) MM001 FIRE peaks, and (**h**) CCS15 FIRE peaks. Kendall’s *τ* is reported as a summary statistic of global ordinal association. Pairwise comparisons between adjacent cell-type specificity categories were assessed using Mann–Whitney U tests, with Holm correction applied to account for multiple testing. Effect sizes for these comparisons were quantified using the rank-biserial correlation (*r*_*rb*_). (* *p*_adj_ *<* 0.05, ** *p*_adj_ *<* 0.01, *** *p*_adj_ *<* 0.001).

**Extended Data Figure 5.**
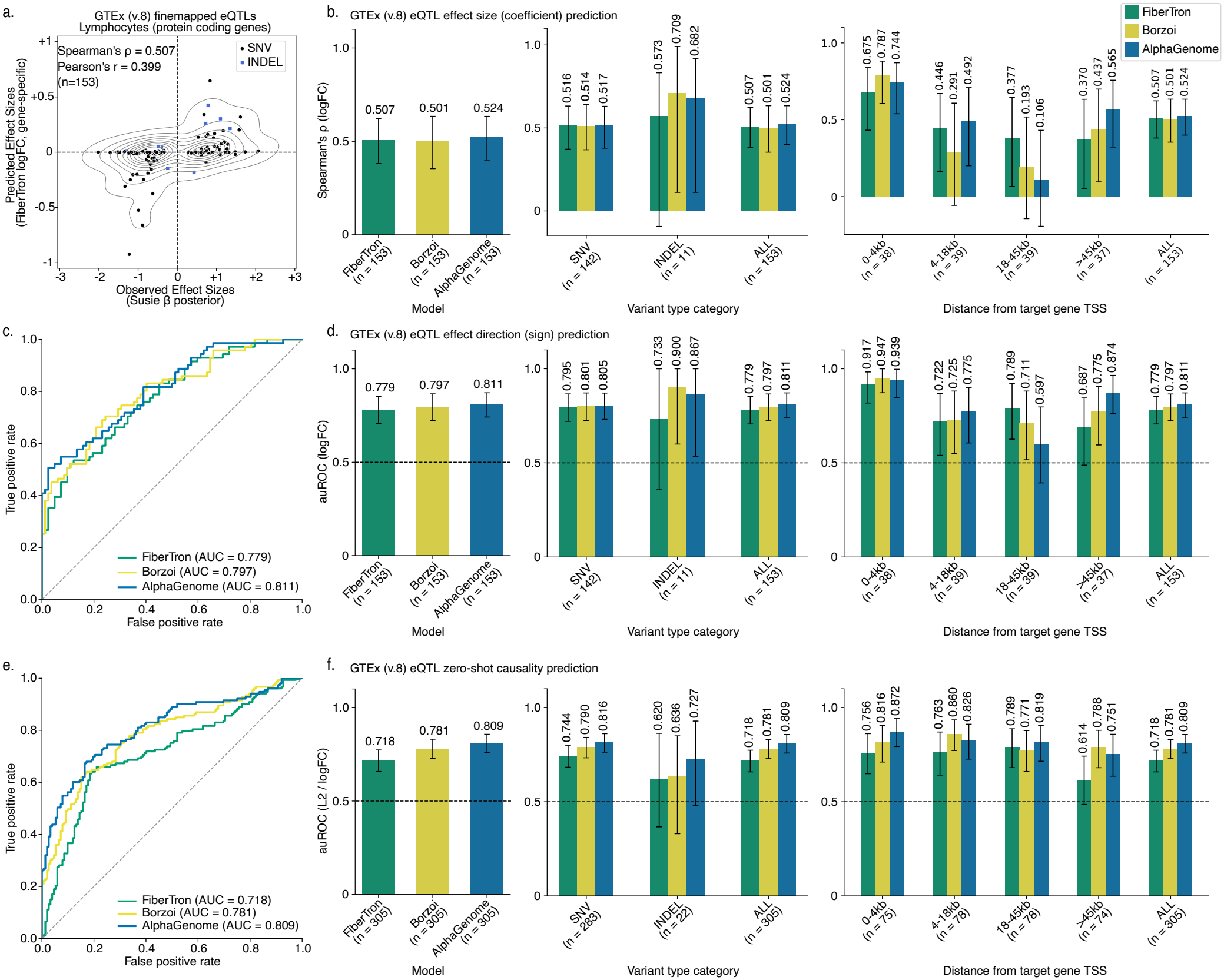
Expression quantitative trait loci (eQTL) prediction tasks. **a**. Comparison of observed effect sizes (SuSiE^109^ *β* posterior estimates) and FiberTron-predicted effect sizes (gene-specific logFC scores). **b-f**. Comparative analysis of performance in eQTL prediction tasks between FiberTron, Borzoi^9^, and AlphaGenome^10^. Across all evaluated tasks, FiberTron achieves performance comparable to that of the heavyweight foundational Borzoi and AlphaGenome models in (**b**) effect size (coefficient) prediction, (**c-d**) effect direction (sign) prediction, and (**e-f**) zero-shot causality prediction. **c** and **e** present auROC curves for the corresponding classification tasks. Performance in **b, d**, and **f** is reported both globally (left) and stratified by variant type (middle) and distance to the target gene transcription start site (right). Error bars represent bootstrapped 95% confidence intervals (CIs). All visualizations and reported FiberTron performance metrics are based on HG002 data. For AlphaGenome, predictions were derived from the GM12878 total RNA-seq track, whereas for Borzoi, predictions were obtained from the blood RNA-seq track.

**Extended Data Figure 6.**
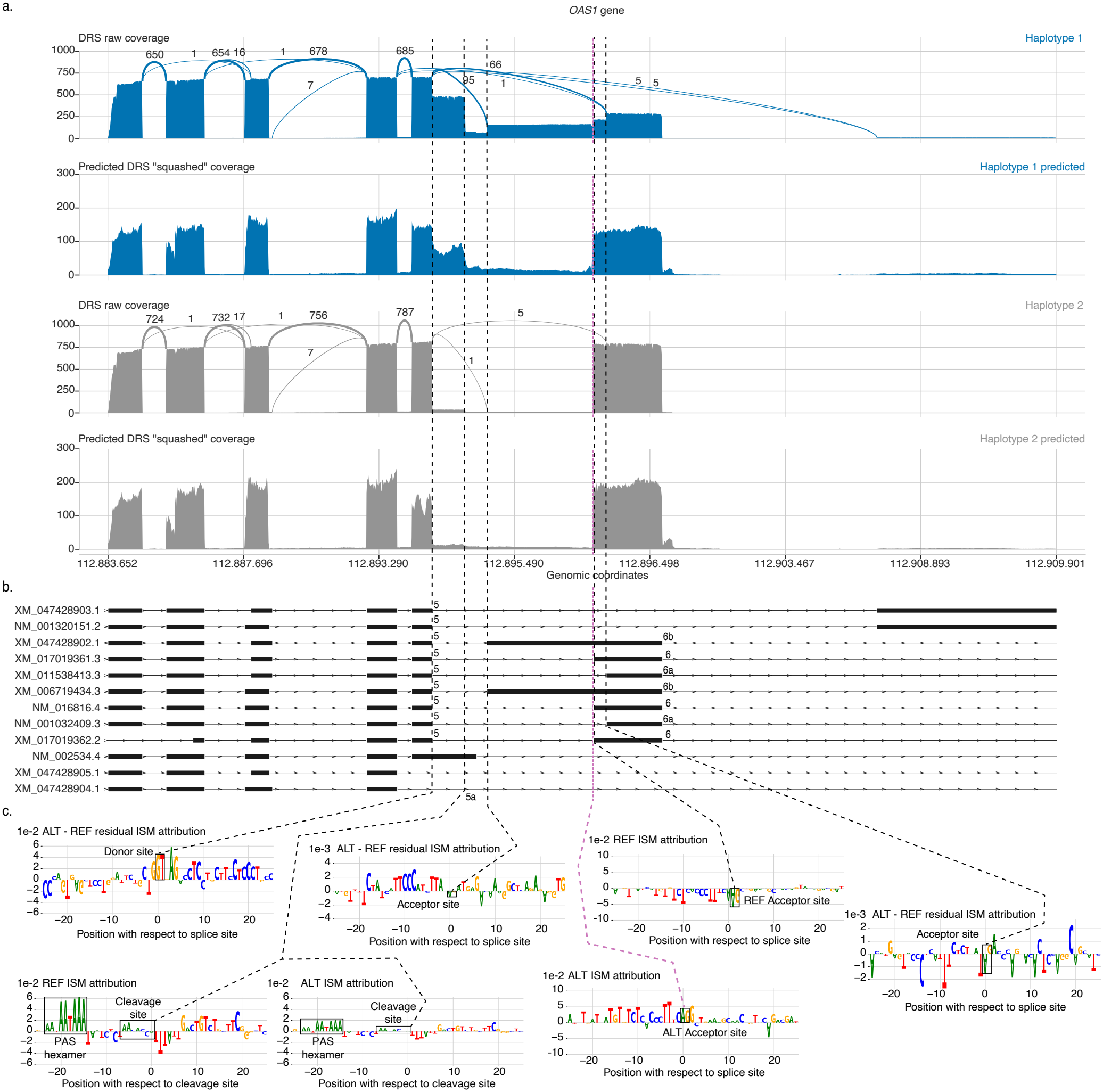
Expanded view of splicing variant-directed *cis*-effects at the *OAS1* locus. **a**. Observed and predicted direct RNA-seq coverage profiles for the reference and alternative alleles across the *OAS1* locus. Observed coverage profiles are shown together with splice junction count data. Putative causal variant chr12:112.896.074:A>G is indicated by the purple dashed line. **b**. Schematic representation of *OAS1* transcript isoforms. **c**. ISM ACWMs at affected splice sites, providing mechanistic insight into the effects of the chr12:112.896.074:A>G variant, including partial intron retention and alternative usage of exons 5 and 6. Exon 5 shows positive residual attribution (REF-ALT) at the donor site (GT). Exon 6 exhibits negative attribution at the reference acceptor site and positive attribution at the alternative acceptor site (AG). Exons 6a and 6b display negative residual attribution at the reference acceptor site (AG). The transcript end site at exon 5a (unannotated transcript) is characterized by a decrease in attribution score for the polyadenylation signal (PAS) hexamer motif (ATTAAA) and cleavage site motifs (CA). All visualizations are based on HG002 data.

**Extended Data Figure 7.**
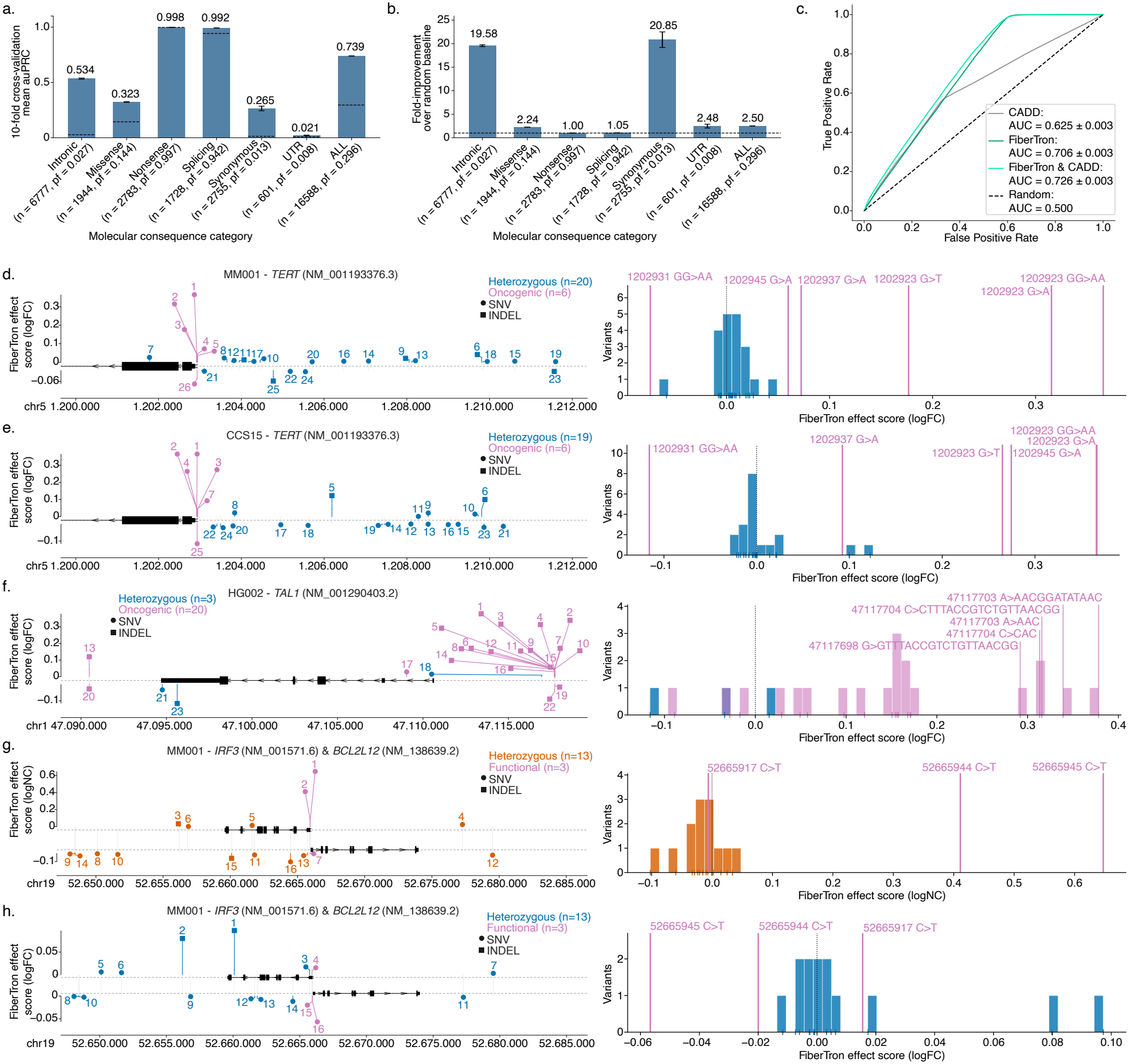
Evaluation of variant pathogenicity prediction and functional effect prediction for disease-associated variants. **a-b**. Supervised pathogenicity classification on the ClinVar^48^ dataset: (**a**) auPRC stratified by molecular consequence category, and (**b**) fold-enrichment in performance relative to a random baseline. **c**. Supervised pathogenicity classification on the gnomAD^49^ dataset. auROC curves for three classifiers trained, respectively, using CADD^50^ scores, FiberTron-derived scores, or their combination as input features. Reported metrics are averages over 10 independent runs on the held-out test set. Visualizations and reported FiberTron performance metrics in **a-c** are based on held-out HG002 data. **d-h**. Plots show gene loci and variant positions. Dots indicate SNVs and squares indicate indels; colors distinguish literature-reported pathogenic/functional variants from within-sample heterozygous variants. Marker height reflects the FiberTron variant effect score, and numbers give each variant’s signed rank by predicted effect. Right panels show the score distributions. In both the (**d**) MM001 cell line and the (**e**) CCS15 patient sample, FiberTron predicts five and four of the six established oncogenic *TERT* promoter mutations to produce the largest increases in gene expression, ranking them among the top positive-effect variants. **(f)** FiberTron also accurately recapitulates the oncogenic activity of clinically relevant neo-enhancer variants near the *TAL1* oncogene^55,56^, predicting increased gene expression associated with these alterations in the HG002 line for 17 of 20 cases. **(g-h)** FiberTron correctly recapitulates the reported functional effects of two out of three non-coding mutations in the bidirectional promoter driving *IRF3* and *BCL2L12* expression^57^, including (**g**) increased CpG methylation and (**h**) a decrease in gene expression. Reported T2T-CHM13 genomic positions chr19:52665917:C>T, chr19:52665944:C>T, and chr19:52665945:C>T correspond to GRCh38 chr19:49665847:C>T, chr19:49665874:C>T, and chr19:49665875:C>T, respectively as reported in Pepe *et al*.^57^.

**Extended Data Figure 8.**
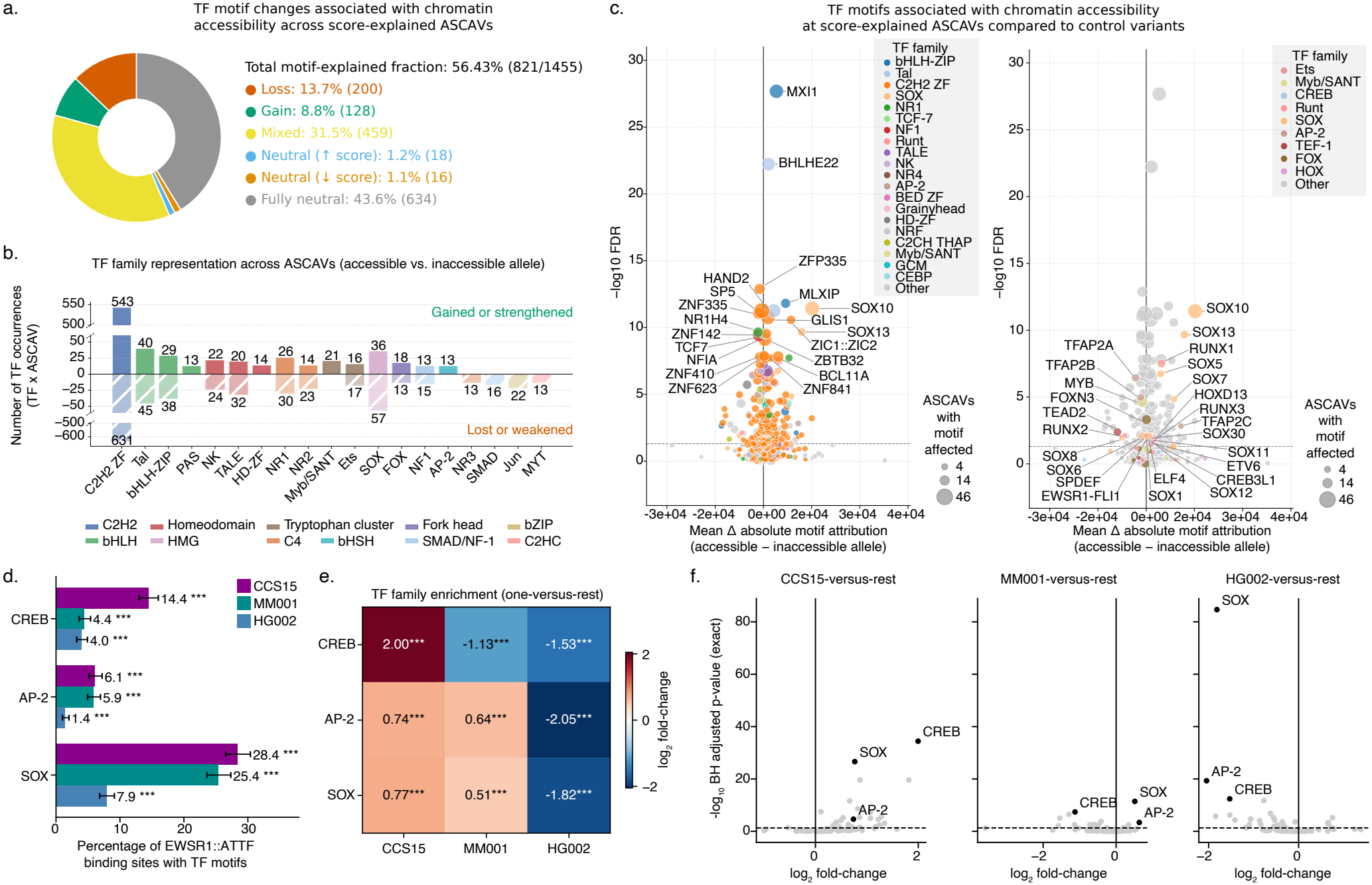
Transcription factor (TF) motif explainability at allele-specific chromatin accessibility variant (ASCAV) sites in melanoma line MM001 and at *EWSR1::ATF1* fusion recruitment sites. **a**. Donut chart depicting the proportion of score-explained ASCAVs that are also explained at the motif level. **b**. TF family representation of motifs affected across explained ASCAV loci, as determined by comparative analysis of accessible versus inaccessible alleles. TF families are color-coded according to their respective class. **c**. Enrichment analysis of affected TF motifs associated with chromatin accessibility at score-explained ASCAV loci compared to control SNVs (one-sided Fischer’s exact test, FDR corrected p-values). In the left volcano plot, the top 20 most significant affected motifs are annotated, whereas the right volcano plot annotates significant affected motifs previously reported in other melanoma lines^6^. TF motif bubbles are color-coded according to their respective TF family. Bubble size represents the number of ASCAVs with the motif affected. **d**. Bar plot summarizing the percentage of EWSR1::ATF1 binding sites containing motifs from the CREB, AP-2, and SOX TF families in CCS15, MM001, and HG002. Error bars indicate 95% Wilson confidence intervals. **e-f**. TF family enrichment analysis at EWSR1::ATF1 binding sites across CCS15, MM001, and HG002. (**e**) Heatmap and (**f**) volcano plots depicting log_2_ fold-changes in TF family motif occurrences between samples. The CREB, AP-2, and SOX families are highlighted in black. Statistical analysis was performed using a paired one-versus-rest exact test with Benjamini–Hochberg (BH) correction for multiple testing. (* *p*_adj_ *<* 0.05, ** *p*_adj_ *<* 0.01, *** *p*_adj_ *<* 0.001).

**Extended Data Figure 9.**
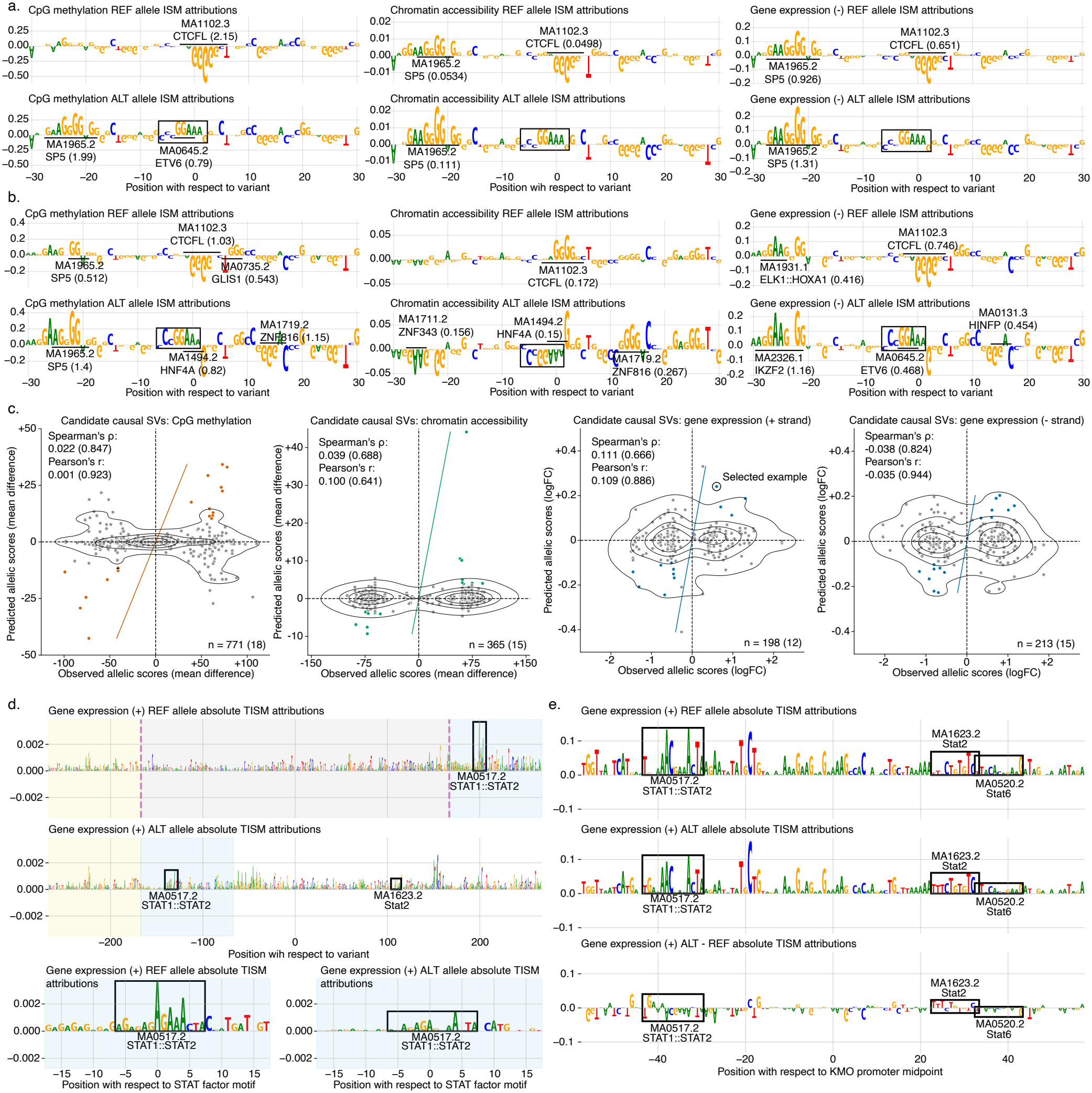
Effect prediction and mechanistic interpretation of complex and disease-associated genetic variants. **a-b**. ISM ACWMs per modality (left to right: CpG methylation, chromatin accessibility, and gene expression) for canonical *TERT* promoter mutation chr5:1.202.923-1.202.924:GG>AA in **(a)** melanoma line MM001 and **(b)** clear cell sarcoma CCS15 patient sample. ETS-like binding motifs (GGAA/T) can be observed for the alternative allele. **c**. Concordance between observed allelic effects and FiberTron-derived structural variant (SV) effect predictions on CpG methylation, chromatin accessibility, and gene expression (plus and minus strand). The concordance of selected candidate causal SVs (colored) is shown in parentheses. An example putative causal variant is circled. **d**. Absolute TISM^34^ ACWMs for the highlighted candidate causal SV. A drop in the attribution score of an identified SV-adjacent STAT-factor motif is observed for the alternative (deleted) allele. **e**. TISM ACWMs in the downstream *KMO* gene promoter show a similar drop in the attribution scores of identified STAT-factor motifs for the alternative allele.

**Extended Data Figure 10.**
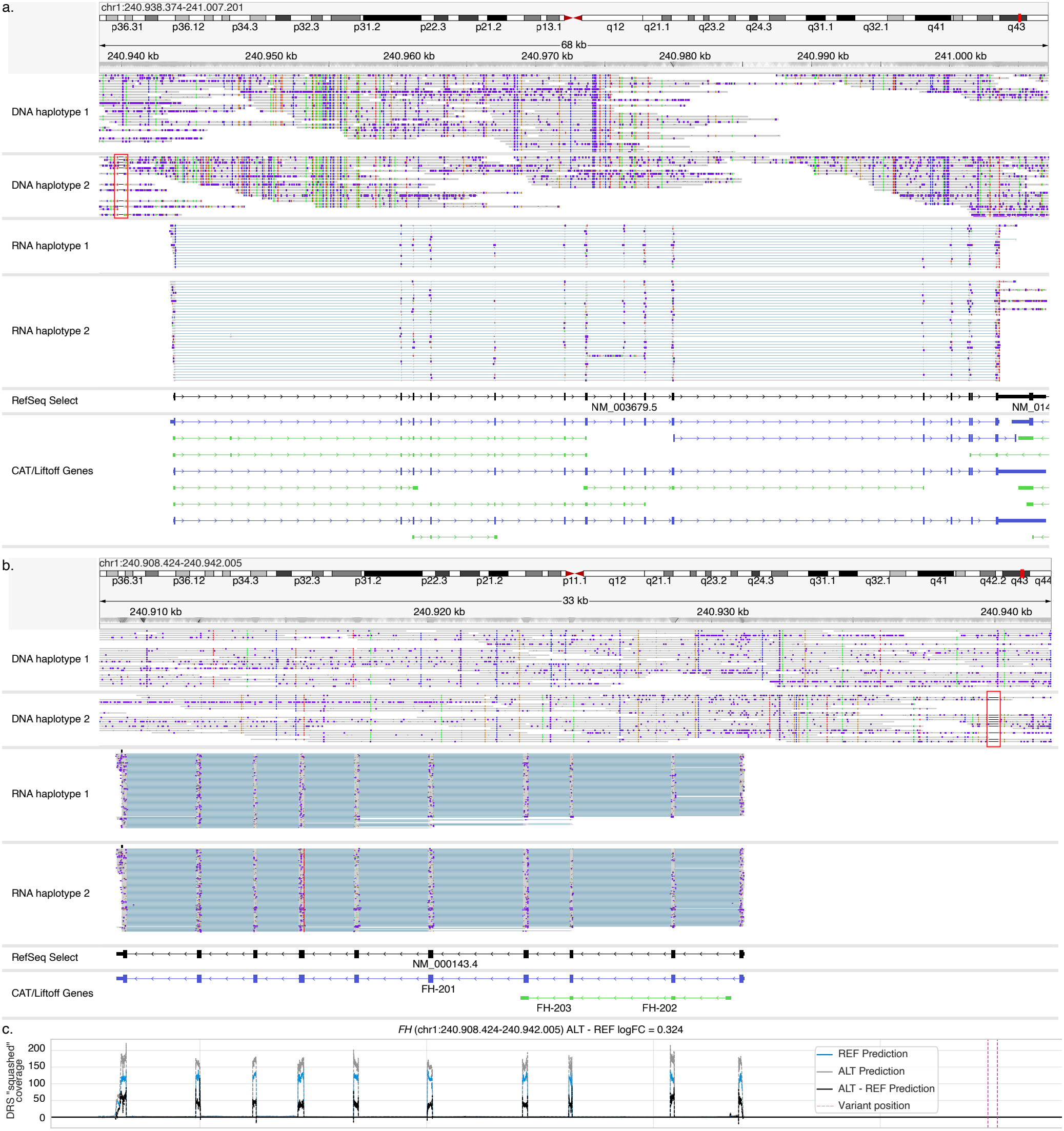
FiberTron predictions and IGV^119^ schematic of the *FH* and *KMO* gene loci for a representative candidate causal SV (334 bp deletion) located at chr1:240.939.801-240.940.135. **a-b**. Haplotype-phased Fiber-seq reads and direct RNA-seq reads showing ASE at the (**a**) *KMO* gene locus (17 versus 36 raw allele-specific counts, one-sided binomial test p-value = 0.00274) and (**b**) *FH* gene locus (328 versus 388 raw allele-specific counts, one-sided binomial test, p-value = 0.0137). The candidate causal SV at chr1:240.939.801-240.940.135 (334 bp deletion) is highlighted by a red box on haplotype 2 in the Fiber-seq alignments. **c**. FiberTron predictions for *FH* gene expression (REF, ALT and ALT - REF), with the candidate causal SV at chr1:240.939.801-240.940.135 (334 bp deletion) denoted with purple dashed lines.

## Supplementary Information

**Supplementary Figure S1.**
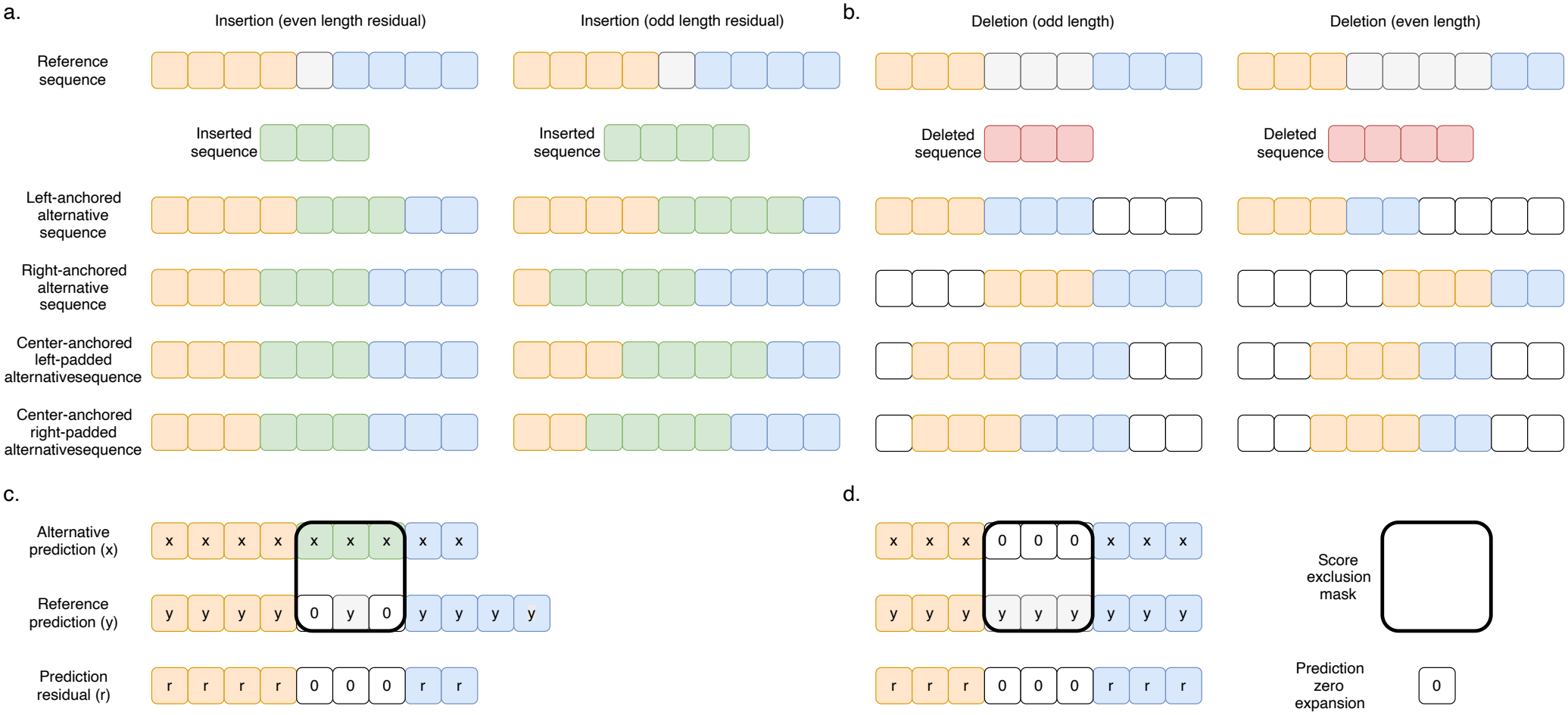
Complex variant effect scoring strategy. **a-b**. Left-anchored, right-anchored, center-anchored left-padded, and center-anchored right-padded alternative sequence generation for (**a**) insertions and (**b**) deletions. **c-d**. Using the left-anchored constructs as representative examples, the prediction re-alignment strategy and exclusion masks used for (**c**) insertion and (**d**) deletion scoring are illustrated.

## Notes

### Competing Interest Statement

The authors have declared no competing interest.

## References

1. Halldorsson, B. V. et al. The sequences of 150,119 genomes in the uk biobank. Nature 607, 732–740 (2022).

2. Wang, X. et al. Deep learning approaches for non-coding genetic variant effect prediction: current progress and future prospects. Briefings Bioinforma. 25, bbae446 (2024).

3. Li, Z. et al. Applications of deep learning in understanding gene regulation. Cell Reports Methods 3 (2023).

4. Avsec, Ž. et al. Base-resolution models of transcription-factor binding reveal soft motif syntax. Nat. genetics 53, 354–366 (2021).

5. Pampari, A. et al. Chrombpnet: bias factorized, base-resolution deep learning models of chromatin accessibility reveal cis-regulatory sequence syntax, transcription factor footprints and regulatory variants. BioRxiv 2024–12 (2025).

6. Atak, Z. K. et al. Interpretation of allele-specific chromatin accessibility using cell state–aware deep learning. Genome research 31, 1082–1096 (2021).

7. Kempynck, N. et al. Crested: modeling genomic and synthetic cell-type-specific enhancers across tissues and species. Nat. Methods 1–14 (2026).

8. Avsec, Ž. et al. Effective gene expression prediction from sequence by integrating long-range interactions. Nat. methods 18, 1196–1203 (2021).

9. Linder, J., Srivastava, D., Yuan, H., Agarwal, V. & Kelley, D. R. Predicting rna-seq coverage from dna sequence as a unifying model of gene regulation. Nat. Genet. 1–13 (2025).

10. Avsec, Ž. et al. Advancing regulatory variant effect prediction with alphagenome. Nature 649, 1206–1218 (2026).

11. Logsdon, G. A., Vollger, M. R. & Eichler, E. E. Long-read human genome sequencing and its applications. Nat. Rev. Genet. 21, 597–614 (2020).

12. Rhie, A. et al. The complete sequence of a human y chromosome. Nature 621, 344–354 (2023).

13. Pardo-Palacios, F. J. et al. Systematic assessment of long-read rna-seq methods for transcript identification and quantification. Nat. methods 21, 1349–1363 (2024).

14. Stergachis, A. B., Debo, B. M., Haugen, E., Churchman, L. S. & Stamatoyannopoulos, J. A. Single-molecule regulatory architectures captured by chromatin fiber sequencing. Science 368, 1449–1454 (2020).

15. Garalde, D. R. et al. Highly parallel direct rna sequencing on an array of nanopores. Nat. methods 15, 201–206 (2018).

16. Peter, C. J. et al. Single chromatin fiber profiling and nucleosome position mapping in the human brain. Cell Reports Methods 4 (2024).

17. Vollger, M. R. et al. Synchronized long-read genome, methylome, epigenome and transcriptome profiling resolve a mendelian condition. Nat. genetics 57, 469–479 (2025).

18. Liu, T. & Conesa, A. Profiling the epigenome using long-read sequencing. Nat. Genet. 57, 27–41 (2025).

19. Dubocanin, D. et al. Integrating single-molecule sequencing and deep learning to predict haplotype-specific 3d chromatin organization in a mendelian condition. bioRxiv (2025).

20. Dixon-Luinenburg, O. et al. Epigenetic conditioning improves sequence-based modeling of gene regulation across cell types and alleles. Biorxiv: Prepr. Serv. for Biol. (2026).

21. Kelley, D. R., Yuan, H., Huang, X. & Linder, J. Cerberus: bidirectional state space blocks improve accuracy and efficiency of regulatory sequence models. bioRxiv 2026–10 (2026).

22. Sasse, A. et al. Benchmarking of deep neural networks for predicting personal gene expression from dna sequence highlights shortcomings. Nat. genetics 55, 2060–2064 (2023).

23. Huang, C. et al. Personal transcriptome variation is poorly explained by current genomic deep learning models. Nat. Genet. 55, 2056–2059 (2023).

24. Barbadilla-Martínez, L., Klaassen, N., van Steensel, B. & de Ridder, J. Predicting gene expression from dna sequence using deep learning models. Nat. Rev. Genet. 1–15 (2025).

25. Dey, K. K. et al. Evaluating the informativeness of deep learning annotations for human complex diseases. Nat. communications 11, 4703 (2020).

26. Ronneberger, O., Fischer, P. & Brox, T. U-net: Convolutional networks for biomedical image segmentation. In Medical image computing and computer-assisted intervention–MICCAI 2015: 18th international conference, Munich, Germany, October 5-9, 2015, proceedings, part III 18, 234–241 (Springer, 2015).

27. Dwarshuis, N. et al. The giab genomic stratifications resource for human reference genomes. Nat. communications 15, 9029 (2024).

28. Kaluscha, S. et al. Evidence that direct inhibition of transcription factor binding is the prevailing mode of gene and repeat repression by dna methylation. Nat. genetics 54, 1895–1906 (2022).

29. Gonzalgo, M. L. & Jones, P. A. Mutagenic and epigenetic effects of dna methylation. Mutat. Res. Mutat. Res. 386, 107–118 (1997).

30. Costello, J. F. et al. Aberrant cpg-island methylation has non-random and tumour-type–specific patterns. Nat. genetics 24, 132–138 (2000).

31. Stefansson, O. A. et al. The correlation between cpg methylation and gene expression is driven by sequence variants. Nat. genetics 56, 1624–1631 (2024).

32. Oxford Nanopore Technologies. modkit: A bioinformatics tool for working with modified bases. https://github.com/nanoporetech/modkit (2024). Accessed: 2026-09-03.

33. Cheetham, S. W., Kindlova, M. & Ewing, A. D. Methylartist: tools for visualizing modified bases from nanopore sequence data. Bioinformatics 38, 3109–3112 (2022).

34. Sasse, A., Chikina, M. & Mostafavi, S. Quick and effective approximation of in silico saturation mutagenesis experiments with first-order taylor expansion. Iscience 27 (2024).

35. Schreiber, J. Tomtom-lite: Accelerating tomtom enables large-scale and real-time motif similarity scoring. Bioinformatics btaf577 (2025).

36. Schreiber, J. tangermeme: A toolkit for understanding cis-regulatory logic using deep learning models. Nat. Methods 1–5 (2026).

37. Castro-Mondragon, J. A. et al. Jaspar 2022: the 9th release of the open-access database of transcription factor binding profiles. Nucleic acids research 50, D165–D173 (2022).

38. Vollger, M. et al. Somatic epimutations cap genetic determinism in the human diploid chromatin epigenome. biorxiv (2024).

39. ENCODE Project Consortium, a. et al. An integrated encyclopedia of dna elements in the human genome. Nature 489, 57 (2012).

40. Degner, J. F. et al. Dnase i sensitivity qtls are a major determinant of human expression variation. Nature 482, 390–394 (2012).

41. DeGorter, M. K. et al. Transcriptomics and chromatin accessibility in multiple african population samples. BioRxiv (2023).

42. Kathail, P. et al. Current genomic deep learning models display decreased performance in cell type-specific accessible regions. Genome Biol. 25, 202 (2024).

43. Hnisz, D. et al. Super-enhancers in the control of cell identity and disease. Cell 155, 934–947 (2013).

44. Wang, Q. S. et al. Leveraging supervised learning for functionally informed fine-mapping of cis-eqtls identifies an additional 20,913 putative causal eqtls. Nat. Commun. 12, 3394 (2021).

45. Quinones-Valdez, G., Amoah, K. & Xiao, X. Long-read rna-seq demarcates cis-and trans-directed alternative rna splicing. Nat. Commun. 16, 9603 (2025).

46. Bonnevie-Nielsen, V. et al. Variation in antiviral 2′,5′-oligoadenylate synthetase (2′5′AS) enzyme activity is controlled by a single-nucleotide polymorphism at a splice-acceptor site in the OAS1 gene. The Am. J. Hum. Genet. 76, 623–633 (2005).

47. Huffman, J. E. et al. Multi-ancestry fine mapping implicates oas1 splicing in risk of severe covid-19. Nat. genetics 54, 125–127 (2022).

48. Landrum, M. J. et al. Clinvar: public archive of relationships among sequence variation and human phenotype. Nucleic acids research 42, D980–D985 (2014).

49. Chen, S. et al. A genomic mutational constraint map using variation in 76,156 human genomes. Nature 625, 92–100 (2024).

50. Schubach, M., Maass, T., Nazaretyan, L., Röner, S. & Kircher, M. Cadd v1. 7: using protein language models, regulatory cnns and other nucleotide-level scores to improve genome-wide variant predictions. Nucleic acids research 52, D1143–D1154 (2024).

51. Rheinbay, E. et al. Analyses of non-coding somatic drivers in 2,658 cancer whole genomes. Nature 578, 102–111 (2020).

52. Huang, F. et al. Tert promoter mutations and monoallelic activation of tert in cancer. oncogenesis 4: e176 (2015).

53. Huang, F. W. et al. Highly recurrent tert promoter mutations in human melanoma. Science 339, 957–959 (2013).

54. Forsyth, R. A. et al. Lrsomatic: a highly scalable and robust pipeline for somatic variant calling in long-read sequencing data. bioRxiv 2026–02 (2026).

55. Mansour, M. R. et al. An oncogenic super-enhancer formed through somatic mutation of a noncoding intergenic element. Science 346, 1373–1377 (2014).

56. Smith, C. et al. Tal1 activation in t-cell acute lymphoblastic leukemia: a novel oncogenic 3’neo-enhancer. Haematologica 108, 1259 (2023).

57. Pepe, D. et al. Reannotation of cancer mutations based on expressed rna transcripts reveals functional non-coding mutations in melanoma. The Am. J. Hum. Genet. 112, 1447–1467 (2025).

58. Li, B. X. & Piesner, J. Establishing preclinical models for clear cell sarcoma of soft tissue. Front. Oncol. 15, 1589773 (2025).

59. Möller, E. et al. Ewsr1-atf1 dependent 3d connectivity regulates oncogenic and differentiation programs in clear cell sarcoma. Nat. communications 13, 2267 (2022).

60. Feng, C. et al. Knocktf: a comprehensive human gene expression profile database with knockdown/knockout of transcription factors. Nucleic acids research 48, D93–D100 (2020).

61. Magnitov, M. D. et al. Mapping functional non-coding variation in individual human genomes through haplotyping, multiomics, and deep learning. Nat. Commun. (2026).

62. Sabbagh, Q., Gilissen, C., Yntema, H. G., Vissers, L. E. & Hoischen, A. Near-perfect genome sequencing in medical genetics. Nat. Genet. 1–10 (2026).

63. Tu, X. et al. Deep genomic models of allele-specific measurements. Genome Res. (2026).

64. Nagai, M., Murphy, A. E., Rizzo, K. & Koo, P. K. Toward generalizable and interpretable ai in regulatory genomics. Nat. Genet. 1–16 (2026).

65. Cheng, J. et al. Alphagenome atlas: in silico mutagenesis of the entire human genome improves prioritization and interpretation of non-coding variants. medRxiv 2026–09 (2026).

66. Swanson, E. G. et al. Mapping single-cell diploid chromatin fiber architectures using daf-seq. Nat. biotechnology 1–12 (2025).

67. Dennis, A. F., Xu, Z. & Clark, D. J. Examining chromatin heterogeneity through pacbio long-read sequencing of m. ecogii methylated genomes: an m6a detection efficiency and calling bias correcting pipeline. Nucleic Acids Res. 52, e45–e45 (2024).

68. de Boer, C. G. & Taipale, J. Hold out the genome: a roadmap to solving the cis-regulatory code. Nature 625, 41–50 (2024).

69. Tu, X., Spiro, A., Chikina, M. & Mostafavi, S. A modality gap in personal-genome prediction by sequence-to-function models. bioRxiv 2026–02 (2026).

70. Spiro, A. E. et al. A scalable approach to investigating sequence-to-function predictions from personal genomes. Nat. Methods 23, 1308–1312 (2026).

71. Schwämmle, T. et al. Smfnet reveals principles of combinatorial transcription factor assembly. bioRxiv 2026–10 (2026).

72. Lee, J., Zhang, K., Shurberg, E., Ramani, V. S. & Pollard, K. S. Chai-net learns a regulatory code for chromatin co-accessibility. bioRxiv 2026–10 (2026).

73. Wang, R., Jin, J., Pott, S. & Zhou, J. Predicting single-molecule chromatin distribution from dna sequence. bioRxiv 2026–10 (2026).

74. Baker, T. M. et al. Genome-wide classification of tumor-derived reads from bulk long-read sequencing. bioRxiv (2026).

75. Hwang, S., Lahoti, A., Dao, T. & Gu, A. Hydra: Bidirectional state space models through generalized matrix mixers, 2024. URL https://arxiv.org/abs/2407.09941 (2024).

76. Sigalova, O. M. et al. Modeling cis-regulatory variation in human brain enhancers across a large parkinson’s disease cohort. bioRxiv (2026).

77. Nature. Genome in a bottle—a human dna standard. Nat. Biotechnol. 33, 675 (2015).

78. Di Tommaso, P. et al. Nextflow enables reproducible computational workflows. Nat. biotechnology 35, 316–319 (2017).

79. Jha, A. et al. Dna-m6a calling and integrated long-read epigenetic and genetic analysis with fibertools. Genome Res. 34, 1976–1986 (2024).

80. Li, H. New strategies to improve minimap2 alignment accuracy. Bioinformatics 37, 4572–4574 (2021).

81. Danecek, P. et al. Twelve years of samtools and bcftools. Gigascience 10, giab008 (2021).

82. Oxford Nanopore Technologies. Dorado: High-performance basecalling for oxford nanopore reads.

83. Ewels, P. A. et al. The nf-core framework for community-curated bioinformatics pipelines. Nat. biotechnology 38, 276–278 (2020).

84. Kovaka, S. et al. Transcriptome assembly from long-read rna-seq alignments with stringtie2. Genome biology 20, 1–13 (2019).

85. Liao, Y., Smyth, G. K. & Shi, W. featurecounts: an efficient general purpose program for assigning sequence reads to genomic features. Bioinformatics 30, 923–930 (2014).

86. Kent, W. J., Zweig, A. S., Barber, G., Hinrichs, A. S. & Karolchik, D. Bigwig and bigbed: enabling browsing of large distributed datasets. Bioinformatics 26, 2204–2207 (2010).

87. Pedersen, B. S. & Quinlan, A. R. Mosdepth: quick coverage calculation for genomes and exomes. Bioinformatics 34, 867–868 (2018).

88. Kelley, D. R. Cross-species regulatory sequence activity prediction. PLoS computational biology 16, e1008050 (2020).

89. Paszke, A. Pytorch: An imperative style, high-performance deep learning library. arXiv preprint arXiv:1912.01703 (2019).

90. Lee, C.-Y., Gallagher, P. W. & Tu, Z. Generalizing pooling functions in convolutional neural networks: Mixed, gated, and tree. In Artificial intelligence and statistics, 464–472 (PMLR, 2016).

91. Zhang, B. & Sennrich, R. Root mean square layer normalization. Adv. Neural Inf. Process. Syst. 32 (2019).

92. Hendrycks, D. & Gimpel, K. Gaussian error linear units (gelus). arXiv preprint arXiv:1606.08415 (2016).

93. Vaswani, A. et al. Attention is all you need. Adv. neural information processing systems 30 (2017).

94. Su, J. et al. Roformer: Enhanced transformer with rotary position embedding. Neurocomputing 568, 127063 (2024).

95. Dauphin, Y. N., Fan, A., Auli, M. & Grangier, D. Language modeling with gated convolutional networks. In International conference on machine learning, 933–941 (PMLR, 2017).

96. Kendall, A., Gal, Y. & Cipolla, R. Multi-task learning using uncertainty to weigh losses for scene geometry and semantics. In Proceedings of the IEEE conference on computer vision and pattern recognition, 7482–7491 (2018).

97. Loshchilov, I. & Hutter, F. Decoupled weight decay regularization. arXiv preprint arXiv:1711.05101 (2017).

98. Ma, J. & Yarats, D. On the adequacy of untuned warmup for adaptive optimization. In Proceedings of the AAAI Conference on Artificial Intelligence, vol. 35, 8828–8836 (2021).

99. Loshchilov, I. & Hutter, F. Sgdr: Stochastic gradient descent with warm restarts. arXiv preprint arXiv:1608.03983 (2016).

100. Lawrence, I. & Lin, K. A concordance correlation coefficient to evaluate reproducibility. Biometrics 255–268 (1989).

101. Auwera, G. A. & O’Connor, B. D. Genomics in the cloud: using docker, gatk, and wdl in terra. (No Title) (2020).

102. Picard toolkit. https://broadinstitute.github.io/picard/ (2019).

103. Zheng, Z. et al. Symphonizing pileup and full-alignment for deep learning-based long-read variant calling. Nat. Comput. Sci. 2, 797–803 (2022).

104. Smolka, M. et al. Detection of mosaic and population-level structural variants with sniffles2. Nat. biotechnology 42, 1571–1580 (2024).

105. Chen, L. et al. Clairs-to: a deep-learning method for long-read tumor-only somatic small variant calling. Nat. Commun. 16, 9630 (2025).

106. Keskus, A. G. et al. Severus detects somatic structural variation and complex rearrangements in cancer genomes using long-read sequencing. Nat. Biotechnol. 44, 247–257 (2026).

107. Lin, J.-H., Chen, L.-C.Yu, S.-C. & Huang, Y.-T. Longphase: an ultra-fast chromosome-scale phasing algorithm for small and large variants. Bioinformatics 38, 1816–1822 (2022).

108. Korsakova, A., Srivastava, D. & Kelley, D. R. Shift augmentation improves dna convolutional neural network indel effect predictions. bioRxiv 2025–04 (2025).

109. Wang, G., Sarkar, A., Carbonetto, P. & Stephens, M. A simple new approach to variable selection in regression, with application to genetic fine mapping. J. Royal Stat. Soc. Ser. B: Stat. Methodol. 82, 1273–1300 (2020).

110. Huang, N. & Li, H. Snp calling, haplotype phasing and allele-specific analysis with long rna-seq reads. Nat. Methods 1–6 (2026).

111. Cheng, J. et al. Accurate proteome-wide missense variant effect prediction with alphamissense. Science 381, eadg7492 (2023).

112. Pedregosa, F. et al. Scikit-learn: Machine learning in python. J. Mach. Learn. Res. 12, 2825–2830 (2011).

113. Majdandzic, A., Rajesh, C. & Koo, P. K. Correcting gradient-based interpretations of deep neural networks for genomics. Genome Biol. 24, 109 (2023).

114. Shrikumar, A., Greenside, P. & Kundaje, A. Learning important features through propagating activation differences. In International conference on machine learning, 3145–3153 (PMlR, 2017).

115. Sundararajan, M., Taly, A. & Yan, Q. Axiomatic attribution for deep networks. In International conference on machine learning, 3319–3328 (PMLR, 2017).

116. Novakovsky, G., Dexter, N., Libbrecht, M. W., Wasserman, W. W. & Mostafavi, S. Obtaining genetics insights from deep learning via explainable artificial intelligence. Nat. Rev. Genet. 24, 125–137 (2023).

117. Simonyan, K., Vedaldi, A. & Zisserman, A. Deep inside convolutional networks: Visualising image classification models and saliency maps. arXiv preprint arXiv:1312.6034 (2013).

118. JGraph. draw.io (2021).

119. Thorvaldsdóttir, H., Robinson, J. T. & Mesirov, J. P. Integrative genomics viewer (igv): high-performance genomics data visualization and exploration. Briefings bioinformatics 14, 178–192 (2013).

